# The clumped isotope signatures of multiple methanogenesis metabolisms

**DOI:** 10.1101/2024.12.18.629299

**Authors:** Jiawen Li, Jeanine L. Ash, Alec Cobban, Briana C. Kubik, Gabriella Rizzo, Mia Thompson, Laetitia Guibourdenche, Stefanie Berger, Kaycee Morra, Ying Lin, Elliott P. Mueller, Andrew L. Masterson, Rebekah Stein, Marilyn Fogel, Mark A. Torres, Xiahong Feng, James F. Holden, Anna Martini, Cornelia U. Welte, Mike Jetten, Edward D. Young, William D. Leavitt

## Abstract

Methane is a potent greenhouse gas, an important energy source, and a potential biosignature on extraterrestrial planetary bodies. The relative abundances of doubly substituted (“clumped”) methane isotopologues (^13^CH_3_D and ^12^CH_2_D_2_) offer important information on the sources and sinks of methane. However, the clumped isotope signatures of microbially produced methane from different methanogenic pathways lack a systematic investigation. In this study, we provide a dataset encompassing the relative isotopologue abundances produced by hydrogenotrophic, methylotrophic, acetoclastic, and methoxydotrophic methanogenesis. We find that a statistical “combinatorial effect” generates significant differences in ^12^CH_2_D_2_ compositions between hydrogenotrophic methanogenesis and other pathways. The thermodynamic drive of methanogenic reactions and phylogenetic affiliation may also influence the isotope compositions of methane. Our study provides new experimental constraints on the isotope signatures of different microbial methanogenic pathways, and evidence of the mechanisms responsible for the observed differences.

**Teaser:** A novel stable isotope tool to track and differentiate sources of biological methane.

## INTRODUCTION

Methane is a critical energy source, greenhouse gas, a key component of the global carbon cycle, and a potential biosignature for life on extraterrestrial planetary bodies. Tracking the sources and sinks of methane is crucial for detecting life beyond Earth (*1, 2*), as well as for reconstructing the methane budget and carbon cycle on Earth (*3*). On Earth, methane is produced by both biological and non-biological processes (*3, 4*). Biogenic methane produced by microbes encompasses a spectrum of biochemical pathways that utilize a variety of substrates. The three major microbial methanogenesis pathways produce methane via hydrogenotrophic (H_2_/CO_2_), methylotrophic (methanol, methylamine, or methyl sulfide, amongst others), or acetoclastic (using acetate) methanogenesis (*5*). More rare yet important biogenic pathways include methoxydotrophy (*6*) and phosphonate demethylation (*7*). These pathways are dominant in different environments and can occur simultaneously in nature (*5, 8–10*). A challenge for quantifying sources and sinks of methane is accurately tracing of the contributions of various metabolic pathways to biogenic methane fluxes in natural environments.

The carbon and hydrogen isotopic ratios of methane molecules offer useful information on the sources of methane. The bulk isotope compositions of methane (^13^C/^12^C and D/H) have been traditionally used to identify its origins (*11, 12*). However, bulk isotope values are influenced by both the isotope compositions of the source materials (e.g., methanol/acetate/CO_2_, H_2_O) that exchange C and H atoms with methane molecules, as well as the isotopic discrimination of carbon and hydrogen during each formation pathway (*13*). More recently, the relative abundance of the doubly-substituted (a.k.a. ‘clumped’) isotopologues of methane (^13^CH_3_D and ^12^CH_2_D_2_) have been applied to identify methane from different sources (*14–16*). Microbial methane produced in laboratory experiments often shows non-equilibrium clumped isotope signatures (*16–19*), whereas microbial methane from some natural environments exhibits isotopologue values closer to thermodynamic equilibrium (*14, 15, 17, 20, 21*). This discrepancy is likely due to the low thermodynamic drive during microbial methanogenesis in some natural settings (e.g., marine sediments, mines), versus the high thermodynamic drive that is common in most laboratory experiments and in some other freshwater or terrestrial settings (*14, 15, 22, 23*).

Despite several pure-culture methanogenesis experiments in which multiply-substituted isotopologues have been measured (*16–19*), there is the need for a better understanding of the effects of different metabolic pathways on isotope clumping. This is very important, both for the interpretation of the sources of methane in nature and for setting endmembers for modeling post-methanogenic alterations (e.g., methane oxidation, mixing, isotopic re-equilibration). Previous studies mainly focus on the fractionation mechanisms of hydrogenotrophic methanogenesis (*22–24*), leaving the other two major biogenic pathways - methylotrophic and acetoclastic - underconstrained. Moreover, a recent report highlights the importance of a so-called ‘combinatorial effect’ on the clumped isotope signatures of methanogenesis using methylated compounds (*19*). In this study, we present a compilation of the isotopic (bulk and clumped) signatures of methanogenic pathways: hydrogenotrophic, methylotrophic, acetoclastic, and methoxydotrophic methanogenesis, with two major goals. First, we investigate if there are differences between the isotope signals of the microbial methanogenic pathways. Second, we elucidate the mechanism for the observed isotopic difference between methanogenic pathways. Furthermore, we highlight the significance of our findings in interpreting the sources of methane in natural environments.

## RESULTS AND DISCUSSION

Our experiments are categorized into nine types based on metabolism and growth conditions (Table 1, Figure 1): H_2_/CO_2_ (35-39 °C), H_2_/CO_2_ (65-80 °C), CH_3_OH + H_2_, CH_3_OH (35-39 °C), CH_3_OH (65 °C), trimethylamine (TMA), TMA + H_2_, 3,4,5-trimethoxybenzoate (TMB), and acetate. The methanogenesis pathways encompass hydrogenotrophic (H_2_/CO_2_), methylotrophic (CH_3_OH, TMA), methoxydotrophic (TMB) and acetoclastic (acetate) methanogenesis. The microbial strains used in this study involve *Methanomassiliicoccus luminyensis* (*M. luminyensis*), *Methermicoccus shengliensis* (*M. shengliensis*), *Methanosarcina mazei* (*M. mazei*), *Methanosarcina barkeri* (*M. barkeri*), *Methanocaldococcus jannaschii* (*M. jannaschii*), *Methanocaldococcus bathoardescens* (*M. bathoardescens*), and *Methanothermococcus thermolithotrophicus* (*M. thermolithotrophicus*). The amount of CH_4_ produced in each culture primarily tracks the amount of substrate provided, as shown in Table 1.

**Fig. 1.**
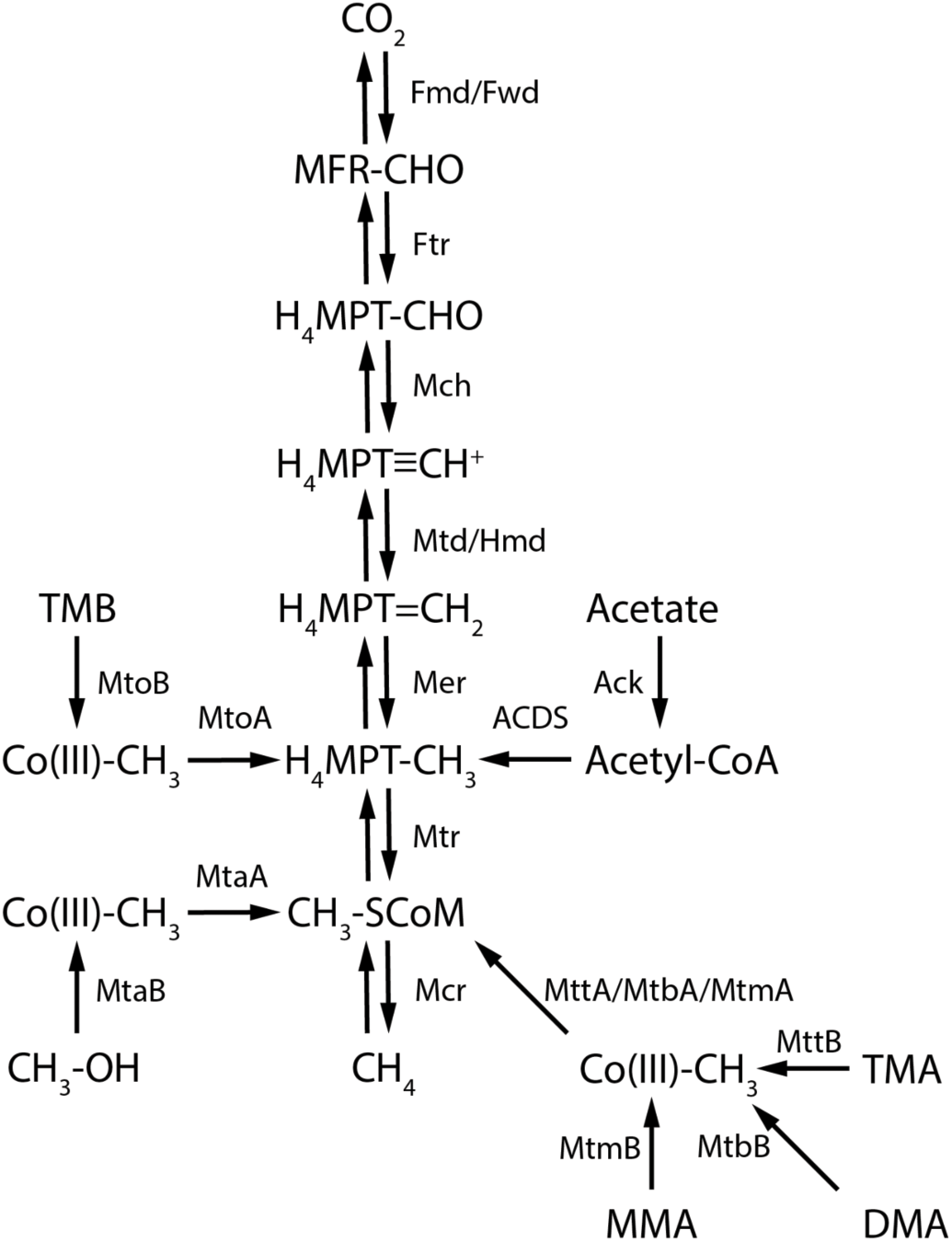
Reaction schemes of the methanogenesis pathways in this study. The reaction scheme is adopted from previous studies (*22, 48, 76, 77*), with the key enzymes and chemical compounds shown on this figure. MFR: methanofuran, H_4_MPT: tetrahydromethanopterin, CoM: coenzyme M, Co(III): cobalamin binding protein, TMB: 3,4,5-trimethoxybenzoate, TMA: trimethylamine, DMA: dimethylamine, MMA: monomethylamine, Fwd/Fmd: formylmethanofuran dehydrogenase, Ftr: formylmethanofurantetrahydromethanopterin formyl-transferase, Mch: methenyltetrahydromethanopterin cyclohydrolase, Mtd: methylenetetrahydromethanopterin reductase, Hmd: H_2_-forming methylenetetrahydromethanopterin dehydrogenase, Mtr: tetrahydromethanopterin S-methyl-transferase, Mcr: methyl-coenzyme M reductase, Ack: acetate kinase, ACDS: Acetyl-CoA decarbonylase/synthase. Mto A-B, Mta A-B, Mtt A-B, Mtb A-B, Mtm A-B are methyltransferases specific to the methylated compounds. For H_2_-dependent methylotrophic methanogenesis (experiments with CH_3_OH + H_2_ and TMA + H_2_ in this study), the microbes lack the enzymes that catalyze the reaction branch between CO_2_ and CH_3_-SCoM (*48*).

**Table 1.** A compilation of isotopic data of lab-cultured methanogenesis experiments in this study and previous studies. All isotopic data are shown in ‰.

| Strain | $\delta D_{H_2O}$ | 1-$\sigma^*$ | Substrate | Temp/$^{\circ}C$ | $\delta^{13}C_{CH_4}$ | 1-σ | $\delta D_{CH_4}$ | 1-σ | $\Delta^{13}CH_3D$ | 1-σ | $\Delta^{12}CH_2D_2$ | 1-σ | total CH<sub>4</sub> (μmol) | Reference |
| --- | --- | --- | --- | --- | --- | --- | --- | --- | --- | --- | --- | --- | --- | --- |
| Rice/Radboud |  |  |  |  |  |  |  |  |  |  |  |  |  |  |
| *M. luminyensis* | -50.3 | 0.1 | CH<sub>3</sub>OH + H<sub>2</sub> | 37 | -97.31 | 0.005 | -271.38 | 0.02 | -2.51 | 0.15 | -44.85 | 0.62 | 1513 | this study |
| *M. luminyensis* | -50.3 | 0.1 | CH<sub>3</sub>OH + H<sub>2</sub> | 37 | -97.96 | 0.004 | -271.95 | 0.01 | -2.15 | 0.15 | -44.5 | 0.58 | 1613 | this study |
| *M. luminyensis* | -50.3 | 0.1 | TMA + H<sub>2</sub> | 37 | -64.91 | 0.003 | -359 | 0.02 | -0.26 | 0.13 | -17.86 | 0.64 | 1222 | this study |
| *M. luminyensis* | -50.3 | 0.1 | TMA + H<sub>2</sub> | 37 | -69.64 | 0.005 | -362.56 | 0.03 | -0.48 | 0.21 | -16.49 | 1.02 | 264 | this study |
| *M. luminyensis* | -50.3 | 0.1 | TMA + H<sub>2</sub> | 37 | -67.95 | 0.005 | -361.87 | 0.03 | 0.08 | 0.18 | -16.55 | 0.94 | 618 | this study |
| *M. shengliensis* | -50.3 | 0.1 | CH<sub>3</sub>OH | 65 | -79.11 | 0.003 | -275.58 | 0.03 | -4.17 | 0.14 | -45.8 | 0.66 | 1671 | this study |
| *M. shengliensis* | -50.3 | 0.1 | CH<sub>3</sub>OH | 65 | -72.15 | 0.003 | -270.38 | 0.03 | -5.51 | 0.12 | -47.82 | 0.56 | 1794 | this study |
| *M. shengliensis* | -50.3 | 0.1 | CH<sub>3</sub>OH | 65 | -66 | 0.005 | -269.49 | 0.03 | -3.22 | 0.21 | -45.7 | 0.8 | 2069 | this study |
| *M. shengliensis* | -50.3 | 0.1 | TMB | 65 | -86.65 | 0.005 | -282.04 | 0.02 | 0.36 | 0.21 | -34.29 | 0.72 | 1064 | this study |
| *M. mazei* | -50.3 | 0.1 | CH<sub>3</sub>OH | 39 | -108.64 | 0.004 | -287.12 | 0.03 | -3.51 | 0.16 | -51.85 | 0.83 | 408 | this study |
| *M. mazei* | -50.3 | 0.1 | CH<sub>3</sub>OH | 39 | -108.73 | 0.006 | -287 | 0.03 | -3.95 | 0.22 | -48.1 | 1.04 | 339 | this study |
| *M. mazei* | -50.3 | 0.1 | TMA | 39 | -89.47 | 0.003 | -379.52 | 0.02 | -3.45 | 0.14 | -31.29 | 0.79 | 1512 | this study |
| Dartmouth (substrate) |  |  |  |  |  |  |  |  |  |  |  |  |  |  |
| *M. barkeri* | -54.7 | 0.2 | H<sub>2</sub>+CO<sub>2</sub> | 35 | -73.15 | 0.004 | -536.75 | 0.03 | -1.98 | 0.22 | -9.88 | 0.98 | 93 | this study |
| *M. barkeri* | -54.7 | 0.2 | H<sub>2</sub>+CO<sub>2</sub> | 35 | -71.83 | 0.005 | -536.88 | 0.003 | -2.51 | 0.22 | -9.86 | 0.96 | 88 | this study |
| *M. barkeri* | -55 | 0.3 | CH<sub>3</sub>OH | 35 | -103.9 | 0.005 | -295.19 | 0.02 | -4.43 | 0.12 | -47.65 | 0.68 | 136 | this study |
| *M. barkeri* | -55 | 0.3 | CH<sub>3</sub>OH | 35 | -98.5 | 0.005 | -290.48 | 0.04 | -9.24 | 0.19 | -47.88 | 0.87 | 185 | this study |
| *M. barkeri* | -55 | 0.3 | CH<sub>3</sub>OH | 35 | -85.93 | 0.005 | -279.4 | 0.02 | -5.21 | 0.22 | -49.48 | 0.75 | 355 | this study |
| *M. barkeri* | -56 | 0.4 | Acetate | 35 | -47.77 | 0.003 | -318.49 | 0.02 | -1.96 | 0.24 | -39.77 | 0.77 | 231 | this study |
| *M. barkeri* | -56 | 0.4 | Acetate | 35 | -48.58 | 0.029 | -319.38 | 0.02 | -1.91 | 0.24 | -41.24 | 0.73 | 223 | this study |
| Dartmouth (water-spike/substrate) |  |  |  |  |  |  |  |  |  |  |  |  |  |  |
| *M. barkeri* | 2937.2 | 3.3 | H<sub>2</sub>+CO<sub>2</sub> | 35 | -75.52 | 0.005 | 806.05 | 0.05 | -4.87 | 0.14 | -16.25 | 0.43 | 81 | this study |
| *M. barkeri* | 2937.2 | 3.3 | H<sub>2</sub>+CO<sub>2</sub> | 35 | -74.6 | 0.012 | 793.24 | 0.04 | -5.27 | 0.14 | -15.51 | 0.45 | 72 | this study |
| *M. barkeri* | 7988 | 6.9 | H<sub>2</sub>+CO<sub>2</sub> | 35 | -74.14 | 0.004 | 3333.25 | 0.18 | -4.57 | 0.13 | -14.6 | 0.31 | 114 | this study |
| *M. barkeri* | 7988 | 6.9 | H<sub>2</sub>+CO<sub>2</sub> | 35 | -78.34 | 0.004 | 3344.29 | 0.17 | -5.03 | 0.11 | -18 | 0.39 | 52 | this study |
| *M. barkeri* | 2491.9 | 2.8 | CH<sub>3</sub>OH | 35 | -95.23 | 0.004 | -60.94 | 0.02 | -10.39 | 0.13 | -10.01 | 0.56 | 167 | this study |
| *M. barkeri* | 2491.9 | 2.8 | CH<sub>3</sub>OH | 35 | -90.26 | 0.003 | -57.29 | 0.02 | -9.99 | 0.31 | -11.61 | 0.57 | 176 | this study |
| *M. barkeri* | 8041.9 | 6.7 | CH<sub>3</sub>OH | 35 | -93.7 | 0.007 | 374.73 | 0.03 | -15.93 | 0.11 | -95.99 | 0.45 | 184 | this study |
| *M. barkeri* | 8041.9 | 6.7 | CH<sub>3</sub>OH | 35 | -79.27 | 0.004 | 402.66 | 0.03 | -15.96 | 0.24 | -68.16 | 0.55 | 352 | this study |
| *M. barkeri* | 2676 | 4.1 | Acetate | 35 | -53.66 | 0.004 | 5.39 | 0.02 | -4.38 | 0.22 | 43.25 | 0.53 | 81.8 | this study |
| *M. barkeri* | 2676 | 4.1 | Acetate | 35 | -53.16 | 0.004 | 0.3 | 0.02 | -4.65 | 0.16 | 39.67 | 0.62 | 106.1 | this study |
| *M. barkeri* | 7909 | 7.9 | Acetate | 35 | -54.09 | 0.004 | 665.18 | 0.05 | -6.41 | 0.33 | 98.67 | 0.61 | 67.1 | this study |
| *M. barkeri* | 7909 | 7.9 | Acetate | 35 | -54.1 | 0.006 | 621.53 | 0.05 | -7.08 | 0.29 | 80.56 | 0.54 | 62.3 | this study |
| U. Mass |  |  |  |  |  |  |  |  |  |  |  |  |  |  |
| *M. jannaschii* | -52.9 | 0.2 | H<sub>2</sub><br>(full)+CO<sub>2</sub> | 80 | -51.93 | 0.009 | -348.6 | 0.03 | 2.75 | 0.3 | -16.28 | 0.82 | 323 | this study |
| *M. bathoardescens* | -53.1 | 0.2 | H<sub>2</sub><br>(full)+CO<sub>2</sub> | 80 | -40.38 | 0.005 | -362.18 | 0.03 | 2.45 | 0.24 | -10.4 | 1.08 | 310 | this study |
| *M. thermolithotrophicus*<sup>†</sup> | -53.5 | 0.1 | H<sub>2</sub><br>(full)+CO<sub>2</sub> | 65 | -39.65 | 0.005 | -403.85 | 0.04 | 0.99 | 0.19 | 1.75 | 0.98 | 376 | this study |
| *M. thermolithotrophicus* | -53.5 | 0.1 | H<sub>2</sub><br>(full)+CO<sub>2</sub> | 65 | -39.22 | 0.004 | -392.46 | 0.03 | 2.67 | 0.13 | -6.27 | 0.84 | 399 | this study |
| *M. thermolithotrophicus* | -53.5 | 0.1 | H<sub>2</sub><br>(full)+CO<sub>2</sub> | 65 | -39.13 | 0.005 | -392.41 | 0.03 | 2.16 | 0.13 | -8.08 | 0.76 | 344 | this study |
| *M. jannaschii* | -52.9 | 0.2 | H<sub>2</sub><br>(full)+CO<sub>2</sub> | 80 | -50.07 | 0.007 | -342.57 | 0.03 | 3.99 | 0.15 | -14.13 | 0.82 | 321 | this study |
| *M. jannaschii* | -53 | 0.2 | H<sub>2</sub> (30 mL)<br>+ CO<sub>2</sub> | 80 | -61.26 | 0.008 | -368.63 | 0.03 | 3.56 | 0.25 | -10.25 | 1.1 | 137 | this study |
| *M. bathoardescens* | -53.4 | 0.1 | H<sub>2</sub> (30 mL)<br>+ CO<sub>2</sub> | 80 | -57.87 | 0.005 | -364.55 | 0.02 | 3.1 | 0.18 | -10.73 | 0.78 | 194 | this study |
| *M. thermolithotrophicus* | -52.6 | 0.3 | H<sub>2</sub> (30 mL)<br>+ CO<sub>2</sub> | 65 | -61.89 | 0.003 | -383.53 | 0.02 | 3.38 | 0.14 | -10.61 | 0.79 | 203 | this study |
| *M. jannaschii* | -53 | 0.2 | H<sub>2</sub> (10 mL)<br>+ CO<sub>2</sub> | 80 | -71.84 | 0.004 | -362.32 | 0.03 | 3.25 | 0.14 | -11 | 0.96 | 63 | this study |
| *M. thermolithotrophicus* | -52.3 | 0.2 | H<sub>2</sub> (10 mL)<br>+ CO<sub>2</sub> | 65 | -66.25 | 0.004 | -374.26 | 0.03 | 3.6 | 0.14 | -9.96 | 0.81 | 77 | this study |
| *M. bathoardescens* | -53.8 | 0.1 | H<sub>2</sub> (10 mL)<br>+ CO<sub>2</sub> | 80 | -65.96 | 0.004 | -363.71 | 0.03 | 2.58 | 0.13 | -10.36 | 0.8 | 64 | this study |
| Previous work |  |  |  |  |  |  |  |  |  |  |  |  |  |  |
| *M. acetivorans* |  |  | CH<sub>3</sub>OH | 30 | -32.77 | 0.007 | -328.41 | 0.02 | -3.88 | 0.2 | -40.86 | 0.36 |  | (16) |
| *M. acetivorans* |  |  | CH<sub>3</sub>OH | 30 | -32.76 | 0.005 | -328.45 | 0.02 | -3.84 | 0.09 | -43.24 | 0.31 |  | (16) |
| *M. barkeri* |  |  | CH<sub>3</sub>OH | 30 | -56.55 | 0.007 | -340.18 | 0.02 | -1.11 | 0.13 | -34.33 | 0.35 |  | (16) |
| *M. thermolithotrophicus* |  |  | H<sub>2</sub>+CO<sub>2</sub> | 65 | -49.36 | 0.011 | -394.37 | 0.02 | 2.66 | 0.1 | -19.44 | 0.3 |  | (16) |
| *M. aeolicus* |  |  | H<sub>2</sub>+CO<sub>2</sub> | 46 | -53.22 | 0.007 | -390.9 | 0.02 | 2.83 | 0.22 | -17.34 | 0.34 |  | (17) |
| *M. aeolicus* |  |  | H<sub>2</sub>+CO<sub>2</sub> | 46 | -51.45 | 0.003 | -399.6 | 0.01 | 3.8 | 0.3 | -16.94 | 0.3 |  | (17) |
| *M. barkeri* |  |  | CH<sub>3</sub>OH | 37 | -53.7 | 0.004 | -250.1 | 0.01 | -2.37 | 0.01 | -54.49 | 0.36 |  | (17) |
| *M. barkeri* |  |  | CH<sub>3</sub>OH | 37 | -53.36 | 0.005 | -249.88 | 0.01 | -2.26 | 0.01 | -55.34 | 0.34 |  | (17) |
| *M. acetivorans* |  |  | CH<sub>3</sub>OH | 28 | -29.16 | 0.005 | -343.22 | 0.11 | -4.2 | 0.28 | -30 | 1.54 |  | (18) |
| *M. acetivorans* |  |  | CH<sub>3</sub>OH | 28 | -88.22 | 0.005 | -372.49 | 0.09 | -6.4 | 0.32 | -38 | 2.61 |  | (18) |
| *M. acetivorans* |  |  | CH<sub>3</sub>OH | 28 | -29.17 | 0.005 | -343.03 | 0.07 | -3.8 | 0.28 | -35 | 1.69 |  | (18) |
| *M. maripaludis* |  |  | H<sub>2</sub> + CO<sub>2</sub> | 37 | -50.53 | 0.003 | -372.56 | 0.02 | 2.3 | 0.12 | -12.11 | 0.39 |  | (16) |
| *P. stutzeri* |  |  | MPn<sup>‡</sup> | 37 | -99.99 | 0.006 | -299.49 | 0.04 | 0.02 | 0.13 | -52.28 | 0.83 |  | (19) |
| *P. stutzeri* |  |  | MPn | 37 | -100.15 | 0.01 | -300.22 | 0.03 | 0.5 | 0.24 | -51.11 | 0.88 |  | (19) |

\* 1-σ error of the measurements.
† The clumped isotope values of this data point are away from the other two replicates under the same condition and are close to the equilibrium values between 400 to 450 °C (Figure 2). This is potentially due to mishandling during the experiment, thus is excluded from the analysis.
‡ MPn: methylphosphonic acid.

The clumped isotope signatures we measure for each pathway fall within the nominal ‘microbial methanogenesis’ field defined by previous studies (*13, 16, 25*). The Δ^13^CH_3_D values in this study range from -9.24 ± 0.19 to 3.99 ± 0.15 ‰ and Δ^12^CH_2_D_2_ values range from -51.85 ± 0.83 to -6.27 ± 0.84 ‰ (Table 1, Figure 2A). While the bulk isotope signatures exhibit large variations among methanogenic pathways, they are generally within the previously-defined range for biogenic methane (*12, 26*). The methane δD ranges from -536.88 ± 0.003 to -269.49 ± 0.03 ‰, while δ^13^C ranges from -108.73 ± 0.006 to -39.13 ± 0.005 ‰ (Table 1, Figure 2B). The isotopic compositions of the samples in this study demonstrate pathway and strain dependence (Table 1, Figure 2), characterized by significant differences in δD and Δ^12^CH_2_D_2_ among different methanogenic pathways (Figure 2).

**Fig. 2.**
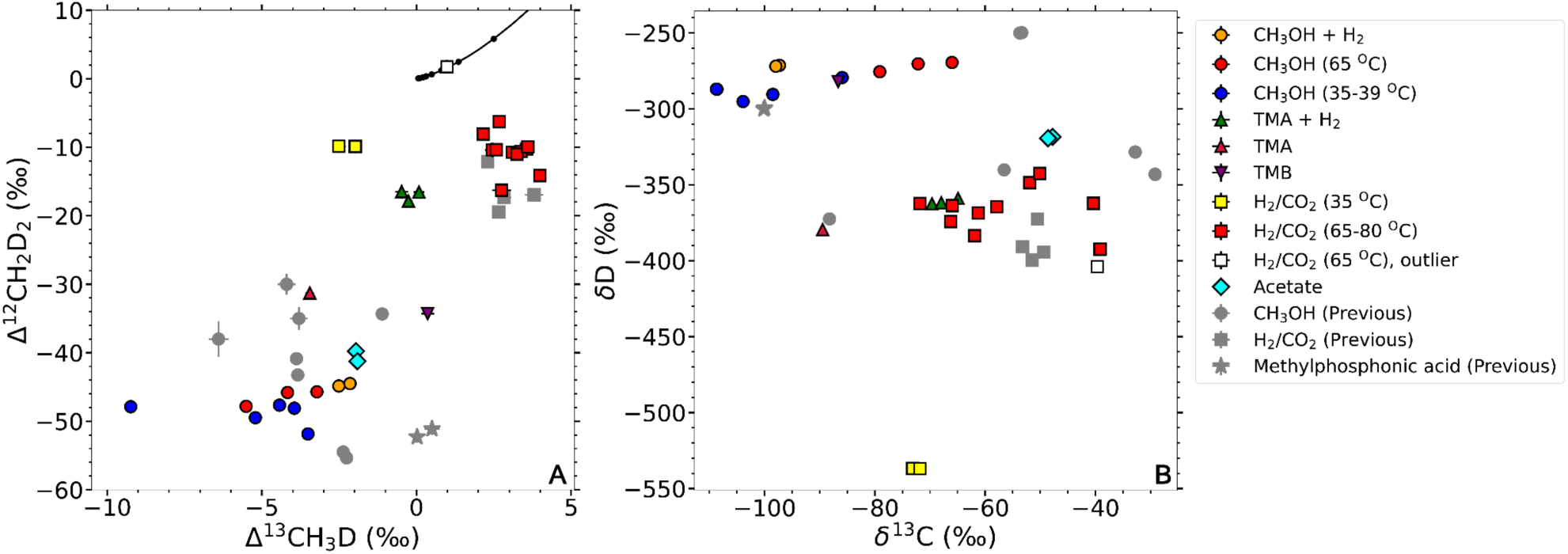
Isotopic data for the non-deuterium-spiked experiments in this study (colored) and previous studies (grey). **(A)** Clumped isotope signatures; **(B)** Bulk isotope signatures. The data used in this figure are shown in Table 1. The solid black line in **panel A** is the thermodynamic equilibrium curve for the doubly-substituted methane isotopologues. All lab-cultured microbial methanogenesis samples included in this figure have both Δ^13^CH_3_D and Δ^12^CH_2_D_2_ values, and show disequilibrium clumped isotope signatures. One biological replicate of hydrogenotrophic methanogenesis by *M. thermolithotrophicus* (white square) with full H_2_ is different from the other two replicates under the same growth condition, therefore it is marked as an outlier.

The apparent hydrogen isotope fractionation factors between methane and water (^D^α_CH4-H2O_) and carbon isotope fractionation factors between methane and source carbon (^13^α_CH4-Carbon_) are shown in Figure 3 and Table S2. The δD values of waters used in the calculations are listed in Table 1, and the isotopic values of other substrates (CO_2_, methanol, acetate, TMA, TMB) are listed in Table S1. The fractionation factors are broadly dependent on the methanogenesis pathway as well. Hydrogenotrophic methanogenesis generates the largest D/H fractionation (^D^α_CH4-H2O_ between 0.49 to 0.69), followed by methylotrophic methanogenesis with TMA and TMA+H_2_ (^D^α_CH4-H2O_ between 0.65 to 0.67). Methoxydotrophic, methylotrophic, and acetoclastic methanogenesis with TMB, CH_3_OH, CH_3_OH+H_2_, and acetate yield smaller D/H fractionations (^D^α_CH4-H2O_ between 0.72 to 0.77). All the hydrogen fractionation factors deviate from the equilibrium values at the incubation temperatures (Figure 3A), suggesting the domination of kinetic isotope effects that are typical in laboratory culture experiments. The available ^13^α_CH4-Carbon_ data demonstrate relatively small variations between different methanogenic pathways, ranging from 0.93 to 0.98 (Table S2, Figure 3B).

**Fig. 3.**
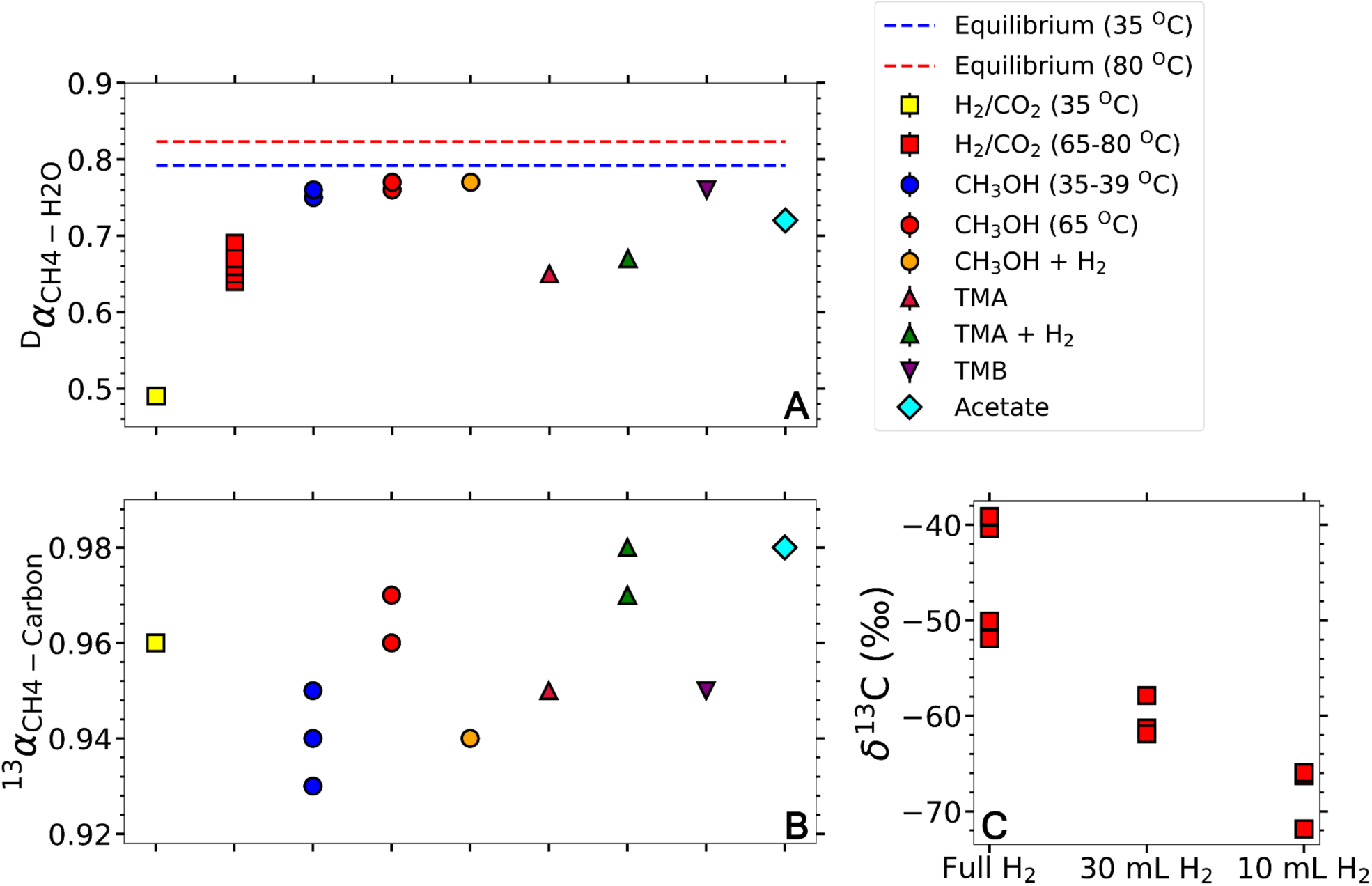
Net carbon and hydrogen fractionation factors with different substrates (Panel A and B), and bulk carbon isotope values of hyperthermophilic hydrogenotrophic methanogenesis under different headspace H_2_ (Panel C). The two dashed lines in panel A show the equilibrium isotope fractionation factors at 35 and 80 ^O^C, based on the equation in the previous study (*78*).

In the following sections, we discuss the patterns of bulk and clumped isotope signatures as a function of microbial methanogenesis pathways, the mechanisms for the observed differences among pathways, and how these may be applied to interpret methane in nature.

### Different methanogenesis pathways yield variations of Δ^12^CH_2_D_2_ due to combinatorial effects

The most distinctive pattern is the difference in Δ^12^CH_2_D_2_ values between hydrogenotrophic and other methanogenic pathways (Figure 2A), and it is due to a so-called ‘combinatorial effect’. It follows from the calculation of Δ^12^CH_2_D_2_ in the stochastic reference frame (*27, 28*). Briefly, because the stochastic distribution of isotopologues is calculated based on bulk isotope ratios assuming, by necessity, that the arithmetic mean of the D/H ratios comprising the molecules applies. However, where more than one D/H ratio contributes to the hydrogen pool for the molecules, the appropriate reference to calculate the stochastic distribution would be the geometric mean of these different ratios. The geometric mean will always be less than the arithmetic mean, meaning that the calculated stochastic ratios will be overestimated and the resulting Δ value underestimated. Therefore, a negative shift in Δ^12^CH_2_D_2_ results when hydrogen atoms from two or more sources with different δD combine to form methane molecules. The sources of distinct D/H ratios can be intracellular due to fractionation at each enzymatic step (*13, 24*) or from different external sources of hydrogen. While the effect can alter Δ^12^CH_2_D_2_ values, it has no effect on Δ^13^CH_3_D values. The extent of the shift reflects the offset between the δD values of the two (or more) hydrogen pools that contribute to methane formation, with a larger offset producing a more significant negative shift (*27, 28*).

The microbial metabolism known as hydrogenotrophic methanogenesis synthesizes methane from carbon dioxide and water, using the electrons from molecular hydrogen (H_2_). Compared with other biogenic pathways, hydrogenotrophic methanogenesis generates the most depleted δD_CH4_ values, but the least negative Δ^12^CH_2_D_2_ values (Figure 2). This would be expected, as all hydrogen atoms originate from one reservoir (*29*), in this case water. In addition, the near-zero intercept of the regression line in Figure 4A lends further support, and is consistent with a previous report (*30*). Due to the hydrogen isotope fractionation during each hydrogen addition step or formation of hydrogen-carrying species (e.g., F_420_H_2_ or HS-CoB), there is a difference between the δD values of hydrocarbon compounds and the hydrogen to be added, generating a negative shift in Δ^12^CH_2_D_2_, but the difference is not large enough to produce a significant shift in comparison to other pathways (*16, 22–24*). Therefore, hydrogenotrophic methanogenesis is typically characterized by modest negative Δ^12^CH_2_D_2_ signatures, due to this so-called ‘endogenous combinatorial effect’ (*19*). The dominance of the endogenous combinatorial effect is further confirmed by the deuterium-spiked (D-spiked) hydrogenotrophy experiments, where both Δ^13^CH_3_D and Δ^12^CH_2_D_2_ do not change significantly, while δD_CH4_ covaries with δD_H2O_ (Figure 4).

**Fig. 4.**
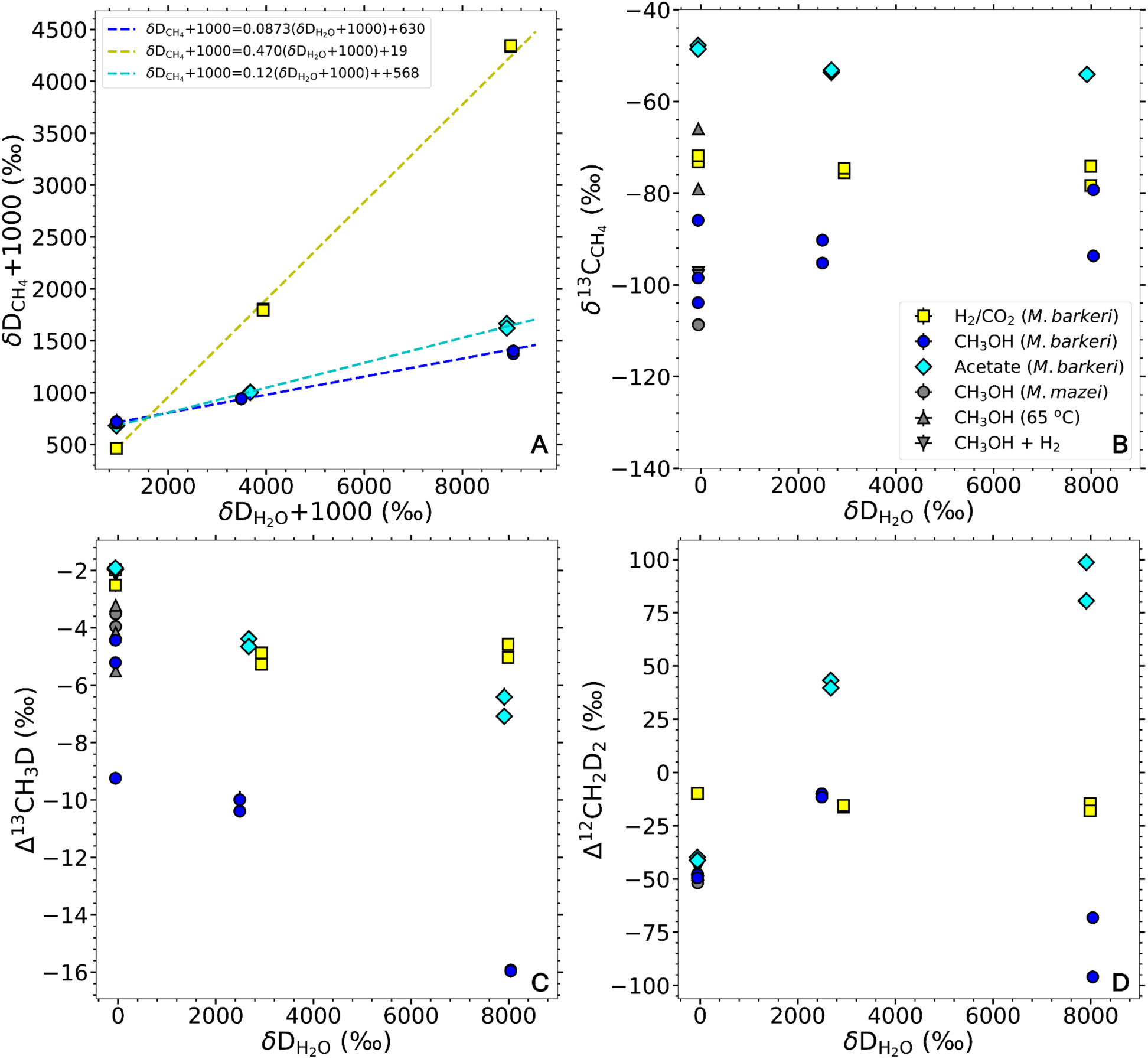
Isotopic signatures of methane versus δD of media water in hydrogenotrophic, methylotrophic and acetoclastic methanogenesis experiments. **(A)** δD_CH4_ + 1000 vs. δD_H2O_ + 1000; **(B)** δ^13^C_CH4_ vs. δD_H2O_; **(C)** Δ^13^CH_3_D vs. δD_H2O_; **(D)** Δ^12^CH_2_D_2_ vs. δD_H2O_. Panel A uses the δD_CH4_ + 1000 vs. δD_H2O_ + 1000 so that the slopes and intercepts reflect ^D^*α*_p_*f* and ^D^*α*_s_(1 – *f*)δD_CH3 + 1000_ in eqn. 7, while panels B-D show the isotopic values vs. δD_H2O_. The data from D-spiked experiments at Dartmouth College are shown as colored points. For comparison, the data from non-deuterium-spiked methylotrophic experiments at Radboud are shown as gray points. It is worth noting that the slopes and intercepts for the methylotrophic and acetoclastic methanogenesis in this figure are derived from the measured values, not the ‘pure’ methylotrophic and acetoclastic endmembers after excluding the contribution from hydrogenotrophic methanogenesis (see the Discussion section for details).

In sharp contrast to hydrogenotrophy, methylotrophic, methoxydotrophic, and acetoclastic methanogenesis generate more negative Δ^12^CH_2_D_2_ and less negative δD_CH4_ values (Figure 2). This is due to the ‘exogenous combinatorial effect’ (*19*), where some hydrogen atoms in product methane originate from the reactant’s methyl group, while the rest source from the cellular water. Exogenous combinatorial effects have been observed in microbial methane formation from methylphosphonate (*19*), and during thermogenic methane formation by pyrolysis (*25*). Our D-spiked experiment supports this hypothesis. By varying the δD_H2O_, we artificially control the difference between the methyl group and water and observe that Δ^12^CH_2_D_2_ respond to the difference (Figure 4C, 4D).

With the reaction schemes in Table 2, we reproduce the observed variation of methane isotope signatures with δD_H2O_ in methylotrophic and acetoclastic methanogenesis by tuning the 5 free parameters (Table 3) to best-fit the experimental data (detailed in Model design and implementation in Materials and Methods Section). Gruen and colleagues (*30*) reported that more than one in four hydrogen atoms in methane are sourced from water (i.e. *f* in Table 3 is greater than 0.25), which demonstrates the reversibility of the dehydrogenation steps from CH_3_-SCoM to CO_2_. Previous reports showed that a small portion of methane is produced via the hydrogenotrophic pathway even when *M. barkeri* is cultivated solely on methanol or acetate (*31–33*). In our model, we treat this input from hydrogenotrophic pathway as a form of reversibility during the dehydrogenation. Therefore, we simplify the model by setting the fraction of hydrogen from water to 0.25 (i.e. *f* = 0.25 in Table 3), and assume that the final headspace gas in methylotrophic or acetoclastic methanogenesis experiments is a mixture of methane produced from methanol or acetate and H_2_/CO_2_. This separates the gas into two components - a ‘pure’ methylotrophic or acetoclastic endmember with no isotopic exchange between water and the methyl group, and a ‘pure’ hydrogenotrophic endmember possessing the isotopic values of hydrogenotrophic methanogenesis. The isotopic values of the ‘pure’ methylotrophic methanogenesis are calculated from the measured isotopic values in the methylotrophic, acetoclastic and hydrogenotrophic samples by *M. barkeri*, assuming the proportion of ‘pure’ methylotrophic or acetoclastic methanogenesis in the mixture is *r* (detailed in the Supplementary Materials). Variations of isotope values of the ‘pure’ methylotrophic endmember with *r* are shown in Figure 5, which shows more prominent mixing effects at higher δD_H2O_. Figure 5D and 5H show the modeled parabolas in Δ^12^CH_2_D_2_ vs. δD_H2O_ that are expected for a combinatorial isotopologue effect (*19*).

**Fig. 5.**
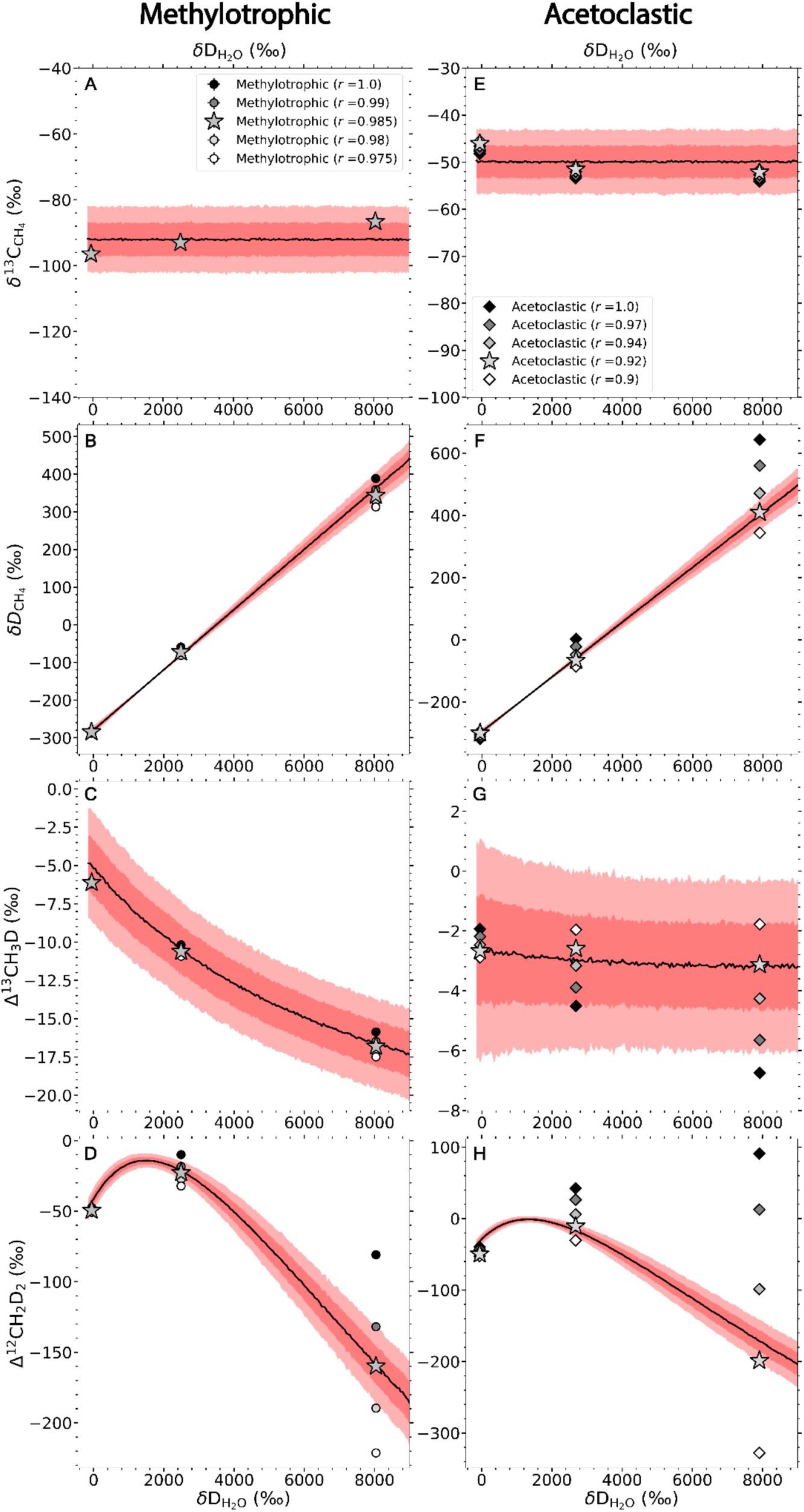
Isotopic models for the D-spiked methylotrophic (panels A-D) and acetoclastic (panels E-H) methanogenesis experiments. The measured signals are assumed to be mixtures of methylotrophic or acetoclastic methanogenesis and hydrogenotrophic methanogenesis. Each panel shows the original measured isotope signatures in black, and deconvolved isotope signatures of the ‘pure’ methylotrophic or acetoclastic endmembers as stars. The mixing ratio *r* denotes the fraction of the methylotrophic or acetoclastic endmember in the mixture. As references, the isotope values derived from a range of *r* values are shown in each panel. The black lines are the mean values from the model, and the deep and light red areas are the 1-σ and 2-σ uncertainty areas, respectively. The reaction schemes, parameters and description of the model are shown in Table 2, Table 3 and the Method Section.

**Table 2.** Reaction schemes and relative reaction rates. ^13^*α*, ^D^*α*_p_, ^D^*α*_s_ are the fractionation factor for ^13^C, and the primary and secondary fractionation factors for hydrogen, respectively. The bracketed variables are the abundances of methane and water isotopologues in the system.

| Reactions* | Reaction rates | Eqn.<br>number |
| --- | --- | --- |
| **For $^{12}\text{C}$ species** |  |  |
| $^{12}\text{CH}_3 - \text{R} + \text{H}_2\text{O} \rightarrow ^{12}\text{CH}_4$ | $r_1 = [^{12}\text{CH}_3\text{R}][\text{H}_2\text{O}]$ | (1) |
| $^{12}\text{CH}_2\text{D} - \text{R} + \text{H}_2\text{O} \rightarrow ^{12}\text{CH}_3\text{D}$ | $r_2 = ^D\alpha_s [^{12}\text{CH}_2\text{DR}][\text{H}_2\text{O}]$ | (2) |
| $^{12}\text{CHD}_2 - \text{R} + \text{H}_2\text{O} \rightarrow ^{12}\text{CH}_2\text{D}_2$ | $r_3 = ^{DD}\gamma_p (^D\alpha_s^2) [^{12}\text{CHD}_2\text{R}][\text{H}_2\text{O}]$ | (3) |
| $^{12}\text{CH}_3 - \text{R} + \text{HDO} \rightarrow ^{12}\text{CH}_4$ | $r_4 = \frac{1}{2} [\text{HDO}][^{12}\text{CH}_3\text{R}]$ | (4) |
| $^{12}\text{CH}_3 - \text{R} + \text{HDO} \rightarrow ^{12}\text{CH}_3\text{D}$ | $r_5 = \frac{1}{2} ^D\alpha_p [\text{HDO}][^{12}\text{CH}_3\text{R}]$ | (5) |
| $^{12}\text{CH}_2\text{D} - \text{R} + \text{HDO} \rightarrow ^{12}\text{CH}_3\text{D}$ | $r_6 = \frac{1}{2} ^D\alpha_s [\text{HDO}][^{12}\text{CH}_2\text{DR}]$ | (6) |
| $^{12}\text{CH}_2\text{D} - \text{R} + \text{HDO} \rightarrow ^{12}\text{CH}_2\text{D}_2$ | $r_7 = \frac{1}{2} (^{DD}\gamma_p) ^D\alpha_p ^D\alpha_s [\text{HDO}][^{12}\text{CH}_2\text{DR}]$ | (7) |
| $^{12}\text{CHD}_2 - \text{R} + \text{HDO} \rightarrow ^{12}\text{CH}_2\text{D}_2$ | $r_8 = \frac{1}{2} (^{DD}\gamma_s) (^D\alpha_s^2) [\text{HDO}][^{12}\text{CHD}_2\text{R}]$ | (8) |
| $^{12}\text{CH}_3 - \text{R} + \text{D}_2\text{O} \rightarrow ^{12}\text{CH}_3\text{D}$ | $r_9 = ^D\alpha_p [\text{D}_2\text{O}][^{12}\text{CH}_3\text{R}]$ | (9) |
| $^{12}\text{CH}_2\text{D} - \text{R} + \text{D}_2\text{O} \rightarrow ^{12}\text{CH}_2\text{D}_2$ | $r_{10} = (^{DD}\gamma_p) ^D\alpha_p ^D\alpha_s [\text{D}_2\text{O}][^{12}\text{CH}_2\text{DR}]$ | (10) |
| **For $^{13}\text{C}$ species** |  |  |
| $^{13}\text{CH}_3 - \text{R} + \text{H}_2\text{O} \rightarrow ^{13}\text{CH}_4$ | $r_{11} = ^{13}\alpha [^{13}\text{CH}_3\text{R}][\text{H}_2\text{O}]$ | (11) |
| $^{13}\text{CH}_2\text{D} - \text{R} + \text{H}_2\text{O} \rightarrow ^{13}\text{CH}_3\text{D}$ | $r_{12} = (^{13}\text{CD}\gamma_s) ^{13}\alpha ^D\alpha_s [^{13}\text{CH}_2\text{DR}][\text{H}_2\text{O}]$ | (12) |
| $^{13}\text{CH}_3 - \text{R} + \text{HDO} \rightarrow ^{13}\text{CH}_4$ | $r_{13} = \frac{1}{2} ^{13}\alpha [\text{HDO}][^{13}\text{CH}_3\text{R}]$ | (13) |
| $^{13}\text{CH}_3 - \text{R} + \text{HDO} \rightarrow ^{13}\text{CH}_3\text{D}$ | $r_{14} = \frac{1}{2} (^{13}\text{CD}\gamma_p) ^{13}\alpha ^D\alpha_p [\text{HDO}][^{13}\text{CH}_3\text{R}]$ | (14) |
| $^{13}\text{CH}_2\text{D} - \text{R} + \text{HDO} \rightarrow ^{13}\text{CH}_3\text{D}$ | $r_{15} = \frac{1}{2} (^{13}\text{CD}\gamma_s) ^{13}\alpha ^D\alpha_s [\text{HDO}][^{13}\text{CH}_2\text{DR}]$ | (15) |
| $^{13}\text{CH}_3 - \text{R} + \text{D}_2\text{O} \rightarrow ^{13}\text{CH}_3\text{D}$ | $r_{16} = (^{13}\text{CD}\gamma_p) ^{13}\alpha ^D\alpha_p [\text{D}_2\text{O}][^{13}\text{CH}_3\text{R}]$ | (16) |

\* In the list of reactions, $\text{CH}_3\text{-R}$ can be either $\text{CH}_3\text{OH}$ or $\text{CH}_3\text{COOH}$ .
† The reaction rates in this table are relative reaction rates, assuming the second-order rate constant for reaction 1 is unity.

**Table 3.** Input parameters of the model for combinatorial effect. The values and uncertainties of the measured parameters are from the experimental data, and those for the fixed parameters are determined by the mixing ratio (*r*) and the fraction of hydrogen from water (*f*), both of which are set at specific values (see the Discussion section for details). The free parameters are assigned to fit the isotopic data of the samples. One set of input parameters is used for each model for the methylotrophic and acetoclastic methanogenesis data, respectively.

| Input parameters | Methylotrophic | 1-σ | Acetoclastic | 1-σ |
| --- | --- | --- | --- | --- |
| **Measured parameters** |  |  |  |  |
| $\delta D_{\text{methyl}}$ (‰) | -54.113 | 0.246 | -159.7 | 1.9 |
| $\delta^{13}C_{\text{methyl}}$ (‰) | -41.343 | 0.017 | -32.2 | 0.05 |
| $\Delta^{13}CH_2D_{\text{methyl}}$ (‰) | +0.842 | 0.352 | 0.0 | 0.300 |
| $\Delta^{12}CD_2H_{\text{methyl}}$ (‰) | +6.255 | 3.776 | 0.0 | 3.000 |
| **Fixed parameters** |  |  |  |  |
| $r$ | 0.985 | - | 0.92 | - |
| $f$ | 0.25 | 0.01 | 0.25 | 0.01 |
| Slope* | 0.0798 | 0.00006 | 0.0879 | 0.00008 |
| Intercept* | 640.373 | 0.090 | 616.925 | 0.097 |
| $^{13}\alpha^{\dagger}$ | 0.9471 | 0.0052 | 0.9817 | 0.0035 |
| $^D\alpha_p^{\ddagger}$ | Slope/$f$ | - | Slope/$f$ | - |
| $^D\alpha_s^{\ddagger}$ | Intercept/$((1-f)(\delta D_{\text{methyl}} + 1000))$ | - | Intercept/$((1-f)(\delta D_{\text{methyl}} + 1000))$ | - |
| **Free parameters** |  |  |  |  |
| $^{13}CD\gamma_p$ | 0.9700 | 0.002 | 0.9960 | 0.002 |
| $^{DD}\gamma_p$ | 0.9700 | 0.002 | 0.9990 | 0.002 |
| $^{13}CD\gamma_s$ | 0.9970 | 0.002 | 0.9975 | 0.002 |
| $^{DD}\gamma_s$ | 0.9950 | 0.002 | 0.9990 | 0.002 |
| **Independent variable** |  |  |  |  |
| $\delta D_{H_2O}$ (‰)<sup>§</sup> | [-150, 9000] | - | [-150, 9000] | - |

\* The values and 1-σ errors of the slopes and intercepts of the regression line are obtained from the least-square regression between $(\delta D_{CH_4} + 1000)$ and $(\delta D_{H_2O} + 1000)$ (eqn. 7, see Materials and Methods for details), following the method in the previous study (79). The regression is conducted on the ‘pure’ methylotrophic and acetoclastic endmember after eliminating the input from hydrogenotrophic methanogenesis (Figure S3).
$^{\dagger} ^{13}\alpha$ is the carbon isotope fractionation factor between methane and substrates, after excluding the input from hydrogenotrophy.
$^{\ddagger} ^D\alpha_p$ and $^D\alpha_s$ are calculated from the slopes, intercepts and $f$ using eqn.7. The uncertainties of $^D\alpha_p$ and $^D\alpha_s$ originate from the uncertainties in the slopes, intercepts and $f$ . They are automatically propagated into the final results.
$^{\S}$ The isotope values are modeled against $\delta D_{H_2O}$ , which ranges from -150 to 9000 ‰, with a 50 ‰ increment (Figure 5).
$^{||}$ The clumped isotope values of the methyl group in acetate are not measured in this study. Therefore, we set the values as 0 and assigned a similar uncertainty as methanol.

The *r* value used in the methylotrophic methanogenesis model is 0.985 (Table 3) -- that is, 98.5 % of total methane comes from methylotrophy, with the remaining from hydrogenotrophy. We choose this value to match the prior observation that 1-2 % of methane comes from hydrogenotrophic methanogenesis during the growth of *M. barkeri* on methanol (*31*). However, the best-fit values for the two primary clumped isotopologue factors (^13CD^γ_p_ and ^DD^γ_p_) significantly deviate from unity (Table 3), demonstrating a large deviation from the rule of geometric mean (*34*). In other words, the fractionation factors of the clumped isotopologues (^13CH3D^*α* and ^12CH2D2^*α*) are smaller than the products of the fractionation factors of the heavy isotopes in the molecules (^13^*α*^D^*α* and ^D^*α*^D^*α*, respectively). However, this is required to fit the isotopic data in this study, based on our analysis of the effects of ^13CD^γ_p_ and ^DD^γ_p_ on the modeled results (Figure S2G and S2L). Notably, we can reproduce the consistent decrease in Δ^13^CH_3_D with increasing δD_H2O_ by applying a smaller ^13CD^γ_p_ (Figure 5C, Figure S2G). Because other studies have generally derived ^13CD^γ_p_ values very near to unity (*15, 22, 23, 30*), we attribute the relatively low values (∼ 0.97) to different enzymes in the methylotrophic methanogenesis pathway. Previous models are based on the reaction schemes of hydrogenotrophic methanogenesis. Methylotrophic methanogenesis, although sharing most of the reaction steps, uses a different set of enzymes to produce CH_3_-SCoM from CH_3_OH (*35, 36*), as shown in Figure 1. Our data imply that the difference in enzymes may yield different net ^13CD^γ_p_ and ^DD^γ_p_ values in methylotrophic methanogenesis.

We rule out several other potential reasons for the observed trend between Δ^13^CH_3_D and δD_H2O_. First, mixing has a minor effect on this trend, as shown in Figure 5C. Second, by stoichiometry, at most 38 % of methanol is consumed throughout the experiment (see the Supplementary Materials), thus a closed-system effect is unlikely to dominate. Moreover, the closed-system effect would drive the net fractionation factors towards unity, which is counter to our observations of the carbon isotope fractionation in methylotrophic methanogenesis (Figure 3, Table S2). Therefore, a large deviation in ^13CD^γ_p_ from unity is the most plausible explanation for these data. It is worth highlighting that the largest δD_H2O_ used in the previous study on the clumped isotope signatures of pure-culture methanogenesis is +335 ‰ (*30*), much smaller than the δD_H2O_ used in this study (Table 1). From our model, a δD_H2O_ of +335 ‰ only produces about a 1 ‰ offset in Δ^13^CH_3_D from lab water with a δD_H2O_ of -50 ‰, which is within the uncertainties between biological replicates.

Acetoclastic methanogenesis is similar to methylotrophic methanogenesis, also with more negative Δ^12^CH_2_D_2_ values (Figure 2). Methane from this pathway is produced by the combination of acetate’s methyl group and hydrogen from an intracellular hydrogen carrier, which is ultimately derived from water (Figure 1). Together, these suggest that the exogenous combinatorial effect occurs in acetoclastic methanogenesis as well. However, unlike methylotrophy, the variation of Δ^12^CH_2_D_2_ with δD_H2O_ in the D-spiked acetoclastic methanogenesis experiments does not yield a simple parabola that is indicative of the exogenous combinatorial effect (Figure 5H), which points to the possibility of a larger input from hydrogenotrophy. To our knowledge, there is no report on the exact proportion of hydrogenotrophic methanogenesis during the cultivation of *M barkeri* on acetate. Therefore, we obtain the *r* value of the acetoclastic methanogenesis experiment from the best fit of experimental data. Since the *f* value in the model is set at 0.25, the best-fit value of the mixing ratio *r* is 0.92 (i.e. 92 % of the total methane comes from acetoclastic methanogenesis). After eliminating the input from hydrogenotrophic methanogenesis, we can obtain a ‘pure’ acetoclastic methanogenesis endmember that does have the characteristic parabola of the exogenous combinatorial effect (Figure 5H). Interestingly, the four clumped isotopologue factors (γ) are much closer to unity, compared with methylotrophic methanogenesis (Table 3). This is consistent with the assumptions and observations in the prior studies that γ’s are close to unity during microbial methanogenesis (*15, 22–24, 30, 37*). Different enzymes used in acetoclastic methanogenesis compared with methylotrophic methanogenesis (Figure 1) could potentially contribute to this difference in γ values.

Methoxydotrophic methanogenesis with TMB also possesses more negative Δ^12^CH_2_D_2_ than hydrogenotrophic methanogenesis, but less negative than acetolactic and methylotrophic methanogenesis. This likely originates from a substantial input of methane (about one third) from carbon dioxide through Wood-Ljungdahl pathway (*6*). This pathway generates acetyl-CoA from carbon dioxide, which is subsequently used to produce methane (*38*). In this process, all hydrogen in the product methane comes from cellular hydrogen (*38*). Therefore, we assume it shows a similar signal as hydrogenotrophic methanogenesis. Since the microbial strain (*M. shengliensis*) used for the methoxydotrophic incubation is a hyperthermophile, we use the average isotope value of hydrogenotrophy under hyperthermophilic conditions as the endmember (Figure 6). Assuming ⅓ of the total methane comes from hydrogenotrophic methanogenesis (*6*), the ‘pure’ methoxydotrophic methanogenesis endmember has a Δ^12^CH_2_D_2_ value of -52.5 ‰, similar to the ‘pure’ methylotrophic and acetoclastic methanogenesis endmembers (Figure 6A, Table S3). Comparing across methylotrophic, acetoclastic and methoxydotrophic methanogenesis, the increasing trend in the Δ^13^CH_3_D values suggests an increasing ^13CD^γ_p_ (Figure 6C, Figure S2G). Again, this pathway dependence of ^13CD^γ_p_ is potentially a result of different enzymes that catalyze the conversion from substrates to CH_3_-SCoM or CH_3_-H_4_MPT (Figure 1).

**Fig. 6.**
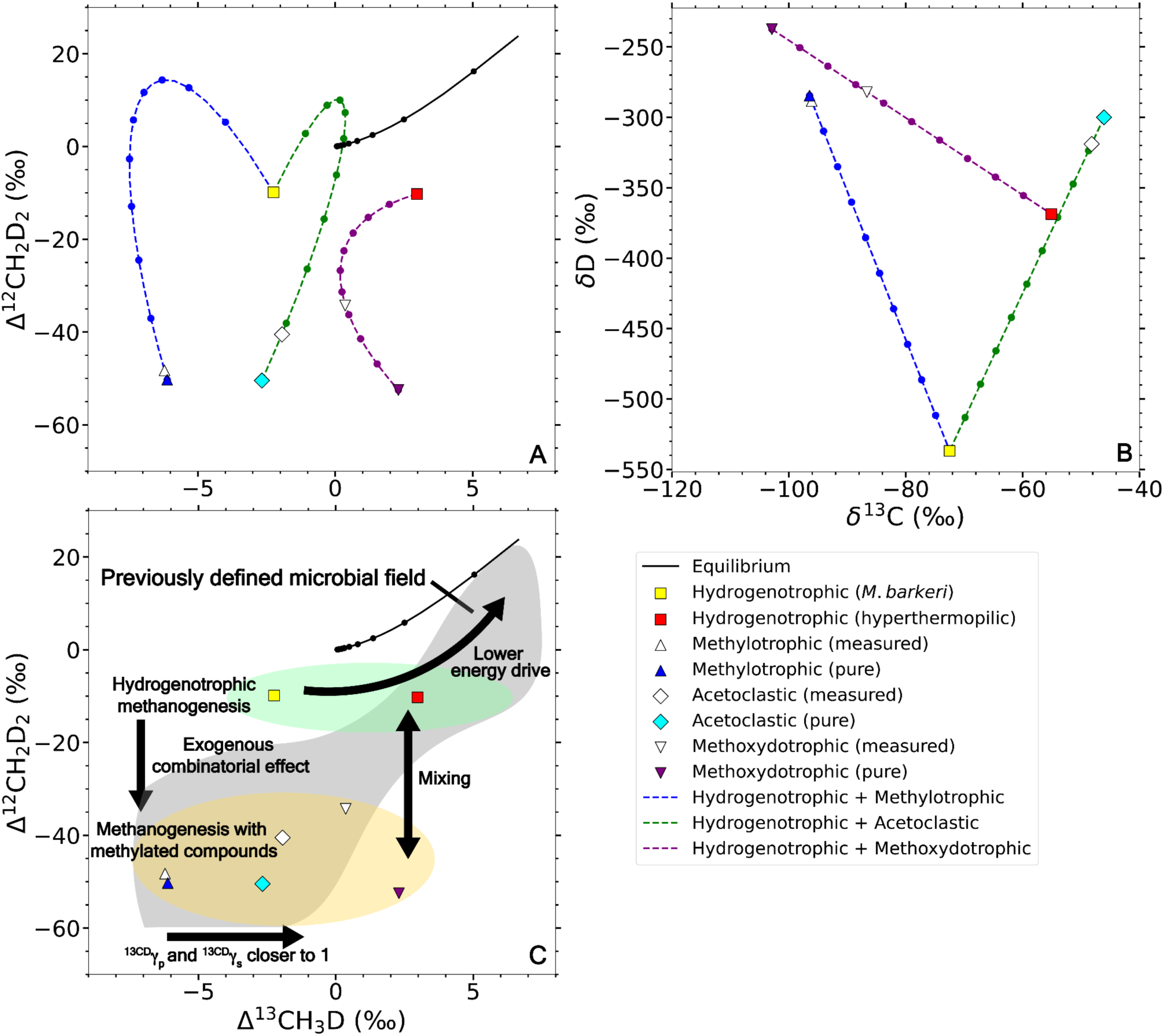
Mixing curves in (A) clumped and (B) bulk isotope spaces, and (C) a summary of the influencing factors on the isotope values of methanogenesis. The average ‘pure’ hydrogenotrophic, methylotrophic, acetoclastic and methoxydotrophic methanogenesis endmembers derived from the mixing model and the corresponding mixing curves are shown in color on panels A and B. The hollowed points are the average measured isotopic values in the experiments. The arrows in Panel C show the processes that can influence the clumped isotope signatures, and the gray area shows the microbial methanogenesis field defined by the previous studies (*13, 25, 66*).

It is proposed that the thermodynamic drive of the methanogenic reactions creates discrepancy between the isotopic signatures of lab-cultured experiments and environmental samples (*14, 15, 22, 23*). However, this effect has only been investigated in hydrogenotrophic methanogenesis (*22, 23, 39*), not for other methanogenic pathways in this study. Future experiments regarding this effect in methylotrophic, acetoclastic and methoxydotrophic methanogenesis are required to resolve this problem. Second, the degree of anti-clumping from the combinatorial effect depends on the difference in the δD of water and the methylated compounds, which varies in nature depending on the environment. Nonetheless, the difference needs to be two orders of magnitude larger (nearly 3000 ‰, as opposed to -50 ‰) to erase the negative Δ^12^CH_2_D_2_ sourced from the exogenous combinatorial effect (Figure 5D, 5H). Therefore, it is unlikely that the natural variation of the source-material δD can eliminate the difference in Δ^12^CH_2_D_2_ between hydrogenotrophic methanogenesis and methanogenesis with methylated compounds (Figure 6A). The clumped isotope signatures of the methyl group in the source material can influence the clumped isotopic values of product methane as well. The clumped isotopic values of naturally-occurring methylated compounds are currently unknown, and will be important directions for future studies, as discussed in the previous study (*19*).

### Variance in isotope values within hydrogenotrophic methanogens

Within hydrogenotrophic methanogens, isotope signatures are different between mesophiles growing at 35 °C, and hyperthermophiles growing from 65 to 80 °C (Figure 2). The isotopic values from hyperthermophilic strains broadly agree with the previous lab-cultured data (*16–19, 30*), as shown in Figure 2. In contrast, both the δD_CH4_ and Δ^13^CH_3_D values are different between mesophilic and hyperthermophilic strains in this study, despite similar δD values of water (Table 1, Figure 2). Moreover, the net hydrogen fractionation factor ^D^α_CH4-H2O_ for mesophiles is 0.470, as represented by the slope of the regression line in Figure 4A. This value is below the previously reported range (0.56 to 0.86) for microbial hydrogenotrophic methanogenesis (*30, 40–44*). In contrast, the ^D^α_CH4-H2O_ for hyperthermophiles in this study ranges from 0.63 to 0.69 (Figure 3A, Table S2), which falls within the previous range. We offer two possible explanations for this difference.

First, we point out that the thermodynamic drive available to the organisms is controlled by the bioenergetic environment provided by the concentrations of substrate, products and temperature. The net Gibbs free energy (ΔG_r_) set by environmental conditions influence the magnitudes of isotope fractionations within the methanogens (*22, 23*). We estimated the net molar Gibbs free energy yield for the hydrogenotrophic methane-forming reaction in each experiment, based on the initial substrate concentration, the final methane production, and substrate consumption by stoichiometry (detailed in the Supplementary Materials). We used these net free energies as input to a model for isotope fractionations associated with each enzyme-mediated step of methane formation during hydrogenotrophic methanogenesis by Gropp et al. (*22*). The net ΔG_r_ values influence the reversibility at each enzymatic step, and thus both overall fractionation between substrates and product methane, as well as the magnitude of the combinatorial effect attending methane formation. The model predicts that δD_CH4_, Δ^13^CH_3_D, and Δ^12^CH_2_D_2_ values will decrease with increasing thermodynamic drive (more negative net ΔG_r_), while the δ^13^C_CH4_ value increases (*22*). The trends predicted by the model fit our experimental data (Figure S1), though the kinetic isotope effects of hydrogen addition steps retain large uncertainties (*22*). Interestingly, the hydrogen isotope fractionations between water and methane is not reproduced by the fractionation model without further parameterization, where we see an offset of ∼200 ‰ in ^2^ε_CH4-H2O_ between the data by *M. barkeri* and the modeled values (Figure S1). This may be due to isotopic disequilibrium between intracellular H_2_ and H_2_O (*22*). Additionally, within hyperthermophilic strains, the methane is lower in 8^13^C values with less initial headspace hydrogen, or equivalently, less thermodynamic drive (Figure 3C). This is consistent with previous reports (*44–46*), suggesting a larger carbon isotope fractionation between product methane and substrate carbons at lower thermodynamic drive.

Alternately, the key difference in the energy conservation approaches between the mesophilic strain (*M. barkeri*) and hyperthermophilic strains (*M. jannaschii, M. bathoardescens*, and *M. thermolithotrophicus*) may contribute to the different Δ^13^CH_3_D values. Similar observations were reported in a previous study, where *M. barkeri* consistently produced methane with negative Δ^13^CH_3_D at growth temperatures of 21 to 38 °C, whereas hyperthermophilic strains produce methane with positive Δ^13^CH_3_D at 30 to 80 °C (*30*). While all the hydrogenotrophs we grow in this study share the same methanogenic pathway (Figure 1), during methanogenesis by *M. barkeri*, the reduction of ferredoxin is independent from the reduction of heterodisufide compound CoM-S-S-CoB. In contrast, during methanogenesis by the hyperthermophilic strains in this study, these two reactions are coupled by flavin-based electron bifurcation (*47*). Further studies on the isotopic fractionations for these two energy conservation processes, and how these values are conserved (or not) among different strains, as well as the impact of thermodynamic drive on the net fractionation between substrate and product, will allow us to better interpret these observations.

### Variance in isotopic values within methylotrophic methanogenesis

Biologically generated methane from methanol shows no significant difference in clumped isotope composition despite different strains and growth temperatures, and our measurements agree with previous studies (Figure 2). This is likely due to the fact that the organisms in this study (*M. shengliensis, M. mazei* and *M. barkeri*) share the same biochemical pathway to convert methanol to methane (*48*) (Figure 1). One exception is *M. luminyensis* performing methanogenesis with CH_3_OH + H_2_. The strain lacks enzymes to catalyze the upper branch of the reactions from CH_3_-SCoM to CO_2_ (*48*). Therefore, it represents a ‘pure’ methylotrophic methanogenesis endmember without the input from hydrogenotrophy. For the other strains, only a negligible amount of methane is produced from hydrogenotrophy when methanol is the only provided substrate, as shown by the D-spiked experiments (Figure 5A-D). The input of hydrogenotrophic methanogenesis cannot significantly alter the ‘pure’ methylotrophic methanogenesis signal without D-enriched water, thus all methylotrophic methanogenesis experiments with methanol and lab water demonstrate similar isotopic signatures (Figure 2).

In contrast, methanogenesis with TMA and TMA+H_2_ shows large differences in clumped isotope signatures, despite similarities in δD of methane (Figure 2). Methanogenesis with TMA demonstrates similar clumped isotope signatures as methanogenesis with CH_3_OH, whereas methanogenesis using TMA+H_2_ carries similar clumped isotope signatures as hydrogenotrophic methanogenesis (Figure 2A). This is contradictory to the expectation from the methanogenesis reaction schemes. In principle, *M. luminyensis* growing on TMA+H_2_ should only produce methane via methylotrophic methanogenesis (*48*), while *M. mazei* should produce methane both via methylotrophic and hydrogenotrophic pathways on TMA (*8*). The reason behind this contradiction is not immediately clear from the available data in this study and our understanding of biochemistry (Figure 1). It is worth highlighting that the reactions from TMA to CH_3_-SCoM use a different set of methyltransferase (MT) enzymes than the reactions from CH_3_OH to CH_3_-SCoM (*35, 36, 49, 50*). The fact that the conversions from tri-, di- and monomethylamine to CH_3_-SCoM are catalyzed by different sets of MT enzymes (*48, 49, 51–53*) further complicates the analysis. Future experiments with D-spiked water and these substrates are necessary to understand the net primary and secondary isotope fractionation factors and test the effect of different enzymes on the isotope fractionation. Nonetheless, our study is the first report on the clumped isotope values of methylotrophic methanogenesis with TMA and opens more directions for future investigations.

### Implications for differentiating methane sources in natural environments

This study provides experimental constraints on the isotopic signatures of methanogenesis via different pathways and direct evidence of the exogenous combinatorial effect that controls those signatures. This work will aid in identifying the sources of methane in natural and human influenced environments. We estimate the ‘pure’ methylotrophic, acetoclastic and methoxydotrophic endmembers from our natural abundance and D-spiked experiments (Figure 6A and 6B). These results highlight the value of Δ^12^CH_2_D_2_ in distinguishing microbial methanogenesis via H_2_/CO_2_ versus methylated compounds, which stems from the exogenous combinatorial effect (Figure 6C). In many of our experiments, hydrogenotrophic methanogenesis accompanies most of the methane producing metabolisms, due to H_2_ production from the fermentation of methylated compounds. This means that the isotopic signatures of microbial methanogenesis will often be a mixture of at least two methanogenic pathways. In natural environments it has been shown that the dominant methanogenic pathway is determined by temperature (*9, 54*) and substrate availability (*8, 10*). Microbial methane produced under high thermodynamic drive in nature usually demonstrates a clumped isotopic signal of mixing between the methanogenesis with H_2_/CO_2_ and methylated compounds (*17, 20, 55–59*), similar to the area of mixing shown in Figure 6C. Although bulk isotopes, especially δD of methane, also show differences between different methanogenic pathways (Figure 6B), they are dependent on the isotopic values of the source materials (*13*). Therefore, it is crucial to use both the bulk and clumped isotope signatures to more reliably interpret the source of methane gas. Additionally, we observe differences in Δ^13^CH_3_D among methylotrophic, acetoclastic, and methoxydotrophic methanogenesis (Figure 2A, Figure 6A). Based on our model (Figure S2), this can be attributed to differences in the primary and secondary clumped isotopologue factors (Figure 5C and 5G), further demonstrating the complexity of microbial control on the clumped isotope signatures.

To our knowledge, this study is the first report of both Δ^13^CH_3_D and Δ^12^CH_2_D_2_ signatures of pure culture methanogenesis with acetate, TMB and TMA. Acetoclastic methanogenesis is a major contributor of methane production in freshwater sediments and anaerobic digesters (*5, 10*). Although less common than hydrogenotrophic and acetoclastic methanogenesis, methanogenesis with TMB and TMA are also important in the methane cycling of some specific environments, for example, coal seams, hypersaline sediments and guts (*6, 8, 48*).

Our study provides a comprehensive look at the methane isotopologue tracers of microbial methanogenesis endmembers. It significantly expands the parameter space for microbial methanogenesis defined by the previous studies, and we provide a mechanistic explanation for the variation in isotope signatures between different methanogenic pathways (Figure 6C). We demonstrate the role of the combinatorial effect in differentiating the metabolisms of H_2_/CO_2_ versus methyl-containing compounds and see the potential role thermodynamic drive plays in controlling the isotope signatures of different methanogenic pathways. Our study contributes to the development of methane clumped isotopes as tools to trace the provenance of methane gases. Results such as these enhance the utility of rare isotopologues of methane as tools for future studies on the sources and sinks of methane in Earth’s atmosphere, and in extra-terrestrial environments on Mars, Titan, and Enceladus.

## Materials and Methods

### Microbial cultivation and methane production

Three strains of methanogens, *Methanomassiliicoccus luminyensis, Methanosarcina mazei and Methermicoccus shengliensis* (referred to as *M. luminyensis, M. mazei* and *M. shengliensis*, respectively), were cultured in-batch. The medium for culturing of *M. mazei, M. luminyensis,* and *M. shengliensis* were prepared with the reagents documented in the Supplementary Materials. All experiments at Radboud University were performed in quadruplicate 125-mL sterile glass serum bottles closed with red butyl stoppers and aluminum crimp caps. Methane production over time was monitored via gas chromatography (GC), and 1 mL of culture media was passed into fresh, sterile media up to 4 times to ensure that the final methane measured was being produced from the substrate of interest for each experiment. After each experiment, cultures were sacrificed by the injection of 10 mL of 6 M NaOH. Samples were then stored in a cool, dark laboratory cabinet until isotopic measurements.

*Methanosarcina barkeri* strain WWM603 (referred to as *M. barkeri* hereafter) was acquired from the Metcalf lab at the University of Illinois (*60–62*) and grown in HS medium supplemented with either H_2_, methanol, or acetate as electron donors (see the Supplementary Materials). Basal HS media was amended with one of the following - no added source, 125 mM methanol, or 40 mM acetate in the anaerobic hood. Following standard medium preparation, 10 mL of media containing each energy source were aliquoted into 26-mL Balch tubes in sextuplet for each electron donor condition. The headspace of each Balch tube was sparged for 5 minutes with 80:20 (v/v) N_2_:CO_2_ and pressurized to 25 psi before autoclaving. The samples without any energy source were pressurized to 25 psi with 80:20 (v/v) H_2_:CO_2_ after inoculation. Cooled tubes were inoculated with 200 µL of *M. barkeri* culture in the late logarithmic growth phase (OD_600_ = 0.27) from hydrogenotrophic growth with no deuterium spike. *M. barkeri* was also cultivated in deuterium-spiked medium water, where the nominated δD values of water were +3000 or +8000 ‰ (V-SMOW). All experiments were inoculated in quintuplicate with a single abiotic control per condition. The samples were then incubated at 35 °C for 92 days, after which the tubes were killed with 100 μL of 1M HCl, and stored inverted (septa down) at 4 °C until isotopic measurements. The abiotic controls were sampled after incubation to confirm the deuterium content of each media water. The final headspace methane amount in the headspace was quantified via GC, using the method described in the previous literature (*63*).

Monocultures of *Methanocaldococcus bathoardescens* (*M. bathoardescens*), *Methanocaldococcus jannaschii* (*M. jannaschii*) were grown at 80 °C, and *Methanothermococcus thermolithotrophicus* (*M. thermolithotrophicus*) was grown at 65 °C with varying amounts of H_2_ in the headspace, using the method described in the previous literature (*64, 65*). Each 60-mL serum bottle contains 25 mL of DSM 282 growth medium (see the Supplementary Materials). The headspace was filled with gas under one of the three conditions: 1. flushed and topped off with 80:20 (v/v) H_2_:CO_2_ (referred to as full H_2_); 2. flushed and topped off with 80:20 (v/v) N_2_:CO_2_, and then 30 mL of headspace was removed and replaced with 30 mL of 80:20 (v/v) H_2_:CO_2_ (referred to as 30 mL H_2_); 3. flushed and topped off with 80:20 (v/v) N_2_:CO_2_, and then 10 mL of headspace was removed and replaced with 10 mL of 80:20 (v/v) H_2_:CO_2_ (referred to as 10 mL H_2_). Before incubation, an additional 100 kPa of either H_2_:CO_2_ (for full H_2_ condition) or N_2_:CO_2_ (for 30 mL and 10 mL H_2_ conditions) was added to each bottle so that at the start of the experiment, they were overpressurized to 2 atmospheric pressure. For each organism and each growth condition, three biological replicates were made, except for *M. thermolithotrophicus* with 10 mL H_2_, where 4 replicates were made. At the end of each experiment, 0.5 mL of the sample from each bottle were taken for cell counting. The final methane production was quantified following a similar procedure as described for *M. barkeri*.

Methane samples were purified on a vacuum purification system using a GC (SRI 8610C GC-TCD) before being analyzed for clumped methane isotopes. The method follows the protocol described in previous studies (*19, 66*). Depending on the concentration of methane, 2-5 mL (for serum bottles) or 15 mL (for Balch tubes) of the headspace gas sample is passively introduced into the vacuum line with a gas-tight syringe, where it is passed over a U-trap chilled with liquid nitrogen (LN_2_) to remove water, carbon dioxide and other species. The sample is then condensed down onto another U-trap with silica gel with liquid nitrogen to trap methane for 30 minutes, after which the non-condensable gasses in the first U-trap are pumped away. The methane is separated further by being passed over a gas chromatograph (GC) with UHP helium as the carrier gas flowing at 20 mL/min. The GC column was held at 25 °C and has two parts: the first (a 3-meter long, 1/8” outer diameter stainless steel column packed with 5A mol sieve) separates N_2_, Ar, H_2_ and O_2_ from CH_4_ while the second (2-meter long, 1/8” outer diameter stainless steel column packed with HaySep D polymer) separates methane from higher hydrocarbons. The purified methane gas was condensed into a glass finger vial with silica gel, and then introduced into the mass spectrometer.

### Isotope notation

Bulk carbon and hydrogen stable isotope ratios use δ - notation, and they are reported as per mil (‰) differences from the international standards VPDB and VSMOW respectively as

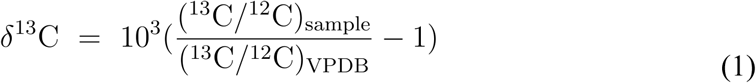

and

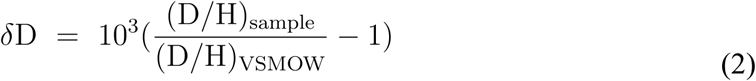

In contrast, the abundances of multiply substituted isotopologues use Δ - notation, and they are reported as per mil differences from a stochastic distribution of isotopologues at infinite temperature as

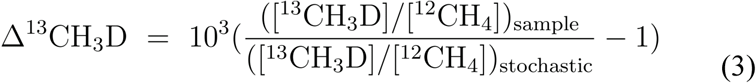

And

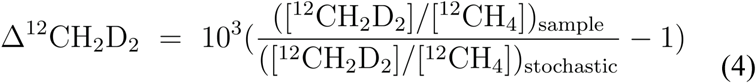

Where the bracketed values are the abundances of the isotopologues.

The isotope fractionation factors in the experiments are expressed as α_A-B_, which denotes the ratio between the heavy to light isotope ratios in two phases A and B:

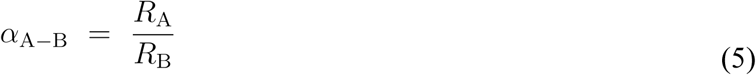

For methane specifically, *R* denotes the ^13^C/^12^C or D/H ratios. To conveniently compare with earlier studies, we also converted α_A-B_ into ε_A-B_:

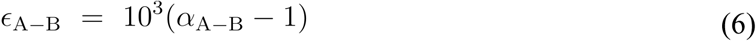

In this study, we focused on the carbon isotope fractionation between methane and the carbon source (CO_2_, methanol, TMA, TMB, acetate), as well as hydrogen isotope fractionation between methane and water.

### Methane isotope ratio measurements

The abundance of two multiply-substituted mass-18 isotopologues of methane (^13^CH_3_D and ^12^CH_2_D_2_) were measured on a Nu Instrument Panorama high-mass-resolution multiple-collector isotope ratio mass spectrometer at the Department of Earth, Planetary, and Space Sciences at the University of California, Los Angeles. The mass spectrometry method follows the application in the previous studies (*19, 59, 66–71*). A cold finger was used to quantitatively remove methane from the sample vial which is heated to ∼60°C to drive gas from the silica gel. Once the gas is transferred, the cold finger and sample bellows are isolated and the cold finger is heated to room temperature. Over 20 minutes, the bellows are expanded and contracted every five minutes to ensure the gas is fully mixed prior to analysis. Two individual sets of analytical methods were used. In the first set, ^12^CH_3_D/^12^CH_4_ and ^12^CH_2_D_2_/^12^CH_4_ were measured as **δ**D and Δ^12^CH_2_D_2_ for 40 blocks of 20 pairs of sample-reference gas measurement cycles. In the second set, ^13^CH_4_/^12^CH_4_ and ^13^CH_3_D/^12^CH_4_ were measured as **δ**^13^C and Δ^13^CH_3_D for 20 blocks of 20 pairs of sample-reference gas measurement cycles. Integration time for each sample or reference gas measurement was 30 seconds. The number of blocks for small-size samples (<100 μmol) were reduced to ensure enough gas for both sets of analytical methods.

Aliquots of Utica gas (a thermogenic gas used as standard at UCLA) were run during and between the analytical sessions. During these sessions, internal precision (2σ) was 0.013, 0.045, 0.378, and 1.308 ‰ for δ^13^C, δD, Δ^13^CH_3_D and Δ^12^CH_2_D_2_ respectively (*n* = 17). The external (95% CI) precision based on repeated runs of Utica gas was determined to be 0.058, 0.160, 0.248, and 0.455 ‰ for δ^13^C, δD, Δ^13^CH_3_D and Δ^12^CH_2_D_2_ respectively (*n* = 17).

### Water and substrate isotope ratio measurements

The δD values of the water for incubations at Dartmouth College were measured at Dartmouth Stable Isotope Laboratory by the method in the previous study (*72*). Water samples that were outside the range of the standards were diluted with a water of known isotopic compositions prior to the measurement using the method described in the previous study (*19*). Water samples were then reduced to molecular hydrogen (H_2_) by hot chromium at 850 °C; the isotope compositions of H_2_ were measured by a dual-inlet isotope ratio mass spectrometer (IRMS, Thermo Delta Plus XL). The final results were reported as δD relative to VSMOW on the VSMOW-SLAP scale with uncertainties <0.5 ‰ (1σ).

Waters used during incubations at Radboud University were measured at Rice University on a Picarro Instruments L2310-*i* cavity ring-down spectrometer with an A0211 vaporization module and attached autosampler to determine δ^18^O and δD. Samples are injected 8 times and run concurrently with 3 in-house standards (Elemental Microanalysis Zero Natural Isotope Water, Elemental Microanalysis Medium Natural Isotope Water, USGS RSIL-W-67400), a drift check of distilled in-house tap water (LT-3) and a quality-control check (USGS45). We report drift and memory corrected averages after van Geldern and Barth (van Geldern and Barth, 2012) of all injections normalized to the VSMOW-SLAP scale. Repeated measurements of USGS45 during this session had an external precision of 0.02 ‰ and 0.07 ‰ (2σ, n = 3) for δ^18^O and δD respectively.

Bulk 8^13^C values of CO_2_ used in the incubations at Dartmouth College were measured following the protocols in a previous study (*73*) using a Gas Bench II coupled to a Delta V Plus isotope ratio mass spectrometer at the Northwestern University Stable Isotope Biogeochemistry Laboratory. Bulk hydrogen δD values were analyzed via a Gas Bench II. Two reference H_2_ gases were standardized separately through repeated analysis of n-alkane reference materials supplied by Indiana University.

The isotopic values of the methyl group in the methanol used in the cultivation at Dartmouth College were analyzed in the Stolper Lab at UC Berkeley, following the previously published method (*74*). Briefly, methanol was reacted in hydriodic acid (HI) to convert methoxyl groups on the methanol to iodomethane (CH_3_I) and then chloromethane (CH_3_Cl) for introduction to the Thermo Scientific 253 Ultra high-resolution dual-inlet isotope-ratio mass spectrometer. Final Δ^13^CH_2_D, and Δ^12^CHD_2_ values are all reported in a stochastic reference frame where 0‰ is equivalent to an infinite temperature for CH_3_Cl, as determined by the previous study (*74*).

Sodium acetate solutions (50 μM in pure methanol) used in the incubations at Dartmouth College were analyzed on a heated electrospray ionization (HESI) Orbitrap QExactive HF (Thermo Fisher, Bremen, Germany) following the published protocol (*75*). This method simultaneously measured the molecular average carbon isotope ratio of acetate and the hydrogen isotope ratio of acetate’s methyl group. Sample isotope ratios were reported on the VPDB and VSMOW scales by comparison to a working standard of sodium acetate (δ^13^C = −19.2‰, δD = −127‰). The solutions were infused into the mass spectrometer using a Vanquish Horizons HPLC Split Sampler Autosampler and a Vanquish Horizons Pump set to 5 μL/min with degassed LC-MS grade methanol as an eluent. An injection volume of 50 μL was carried by the eluent flow to the Orbitrap for a total of 10 acquisition minutes. At that time, the flow rate was increased to 30 μL/min to clear residual sample from the transfer lines. At 10.5 minutes, the flow rate was dropped again to 5 μL/min and 90 seconds later, the next injection began. Data acquisition included all 10 minutes but only integrated between 2 and 8 minutes to calculate isotope ratios. This was repeated to achieve bracketed, sample-standard comparisons (ABABABA, A = standard replicates, B = sample replicates). Acetate standard was diluted to match sample ion current and injected five times. Errors report the standard deviation of the δ^13^C and δD of those replicates.

Bulk carbon and hydrogen isotopes of CH_3_OH, TMA and TMB used during incubations at Radboud University were analyzed at the EDGE Institute at UC Riverside. For δ^13^C measurements, ∼0.7 mg of each TMA/TMB sample and ∼5.6 mg of each CH_3_OH sample was weighed into a low permeability silver capsule and sealed with a liquid seal device then further wrapped in a more permeable tin capsule. These samples were combusted on a Costech ECS 4010 EA connected to a Thermo Scientific Delta V Advantage IRMS via a Conflo IV interface. One gelatin and three USGS glycine standards were analyzed concurrently for data correction. Within-run analytical precision was ∼0.4‰. For δD measurements, ∼1 mg of each sample was weighed into a silver capsule and pyrolyzed on a Thermo Scientific high temperature EA (TC/EA) connected to a Thermo Scientific Delta V Plus IRMS via a Conflo IV interface. Five standards of chicken and turkey feathers, stearic acid, mineral oil, and pump oil were analyzed concurrently to form a regression for data correction. Within-run analytical precision was ∼4‰.

### Model design and implementation

To model the combinatorial effect on the measured clumped isotope signatures, we calculated the abundance of methane isotopologues during methanogenesis using the reaction scheme shown in Table 2. This reaction scheme includes 16 methanogenic reactions between the isotopologues of the methyl group in methanol and the hydrogen atom from water. For simplicity, the reactions that produce triply substituted or heavier clumped isotopologues (e.g., ^12^CHD_3_ and ^13^CH_2_D_2_) are not considered. Since these isotopologues are of very low abundance compared with lighter isotopologues, ruling them out will not cause a significant change in the modeled results for the isotopologues of interest. The relative production rates of methane isotopologues in Table 2 consist of three parts: abundances of isotopologues of the methyl group and water (bracketed values), carbon and hydrogen fractionation factors (^13^*α*, ^D^*α*_p_, and ^D^*α*_s_), and primary and secondary clumped isotopologue factors (^13CD^*γ*_p_, ^13CD^*γ*_s_, ^DD^*γ*_p_, and ^DD^*γ*_s_). Clumped isotopologues factors in methanogenesis are introduced to express the deviation from the rule of geometric mean (*15, 34*). Similar to fractionation factors, the ‘*p*’ and ‘*s*’ in the subscripts of γ represent primary and secondary clumped isotopologue effects, respectively. The relative production rates of isotopologues in Table 2 are treated as equivalent to the relative abundance of isotopologues in the system, which are used to calculate the isotopic values.

The model for the combinatorial effect is applied to the data of D-spiked methylotrophic and acetoclastic experiments at Dartmouth College (Table 1). We categorize the parameters in the model into three parts (Table 3). The measured parameters are determined experimentally, including the isotope values of the substrate methyl groups.

The fixed parameters consist of the fraction of hydrogen from water (*f*), the mixing ratio (*r*), and a series of parameters that depend on these two parameters. In methylotrophic and acetoclastic methanogenesis, the hydrogen atoms in the product methane molecules come from two sources – water and methyl groups of the substrates. The mass balance of hydrogen gives the following equation:

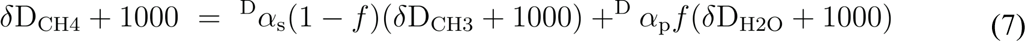

Where ^D^α_p_ and ^D^α_s_ are primary and secondary fractionation factors. Primary fractionation represents the D/H fractionation during the addition of hydrogen to the methyl group, while secondary fractionation represents the D/H fractionation between the methyl group and product methane. *f* is the fraction of hydrogen in methane molecules that comes from water. To simplify the model, we set *f* at 0.25 in the model, meaning one out of four hydrogen atoms in the product methane molecule comes from water. Additionally, we treat the headspace methane as a mixture between hydrogenotrophic and methylotrophic (or acetoclastic) methanogenesis, and denote the portion of methylotrophic or acetoclastic endmembers in the mixture as *r*. Prior to applying the model, the inputs of hydrogenotrophic methanogenesis are eliminated from the measured isotopic values to derive the isotopic values of ‘pure’ methylotrophic and acetoclastic methanogenesis endmembers (detailed calculations of the mixing effect in the Supplementary Materials). We then obtained values of ^D^α_p_*f* and ^D^α_s_(1 – *f*)(δD_CH3_ + 1000) from the slopes and intercepts of the weighted least-square linear regression between (δD_CH4_ + 1000) and (δD_H2O_ + 1000) on the ‘pure’ methylotrophic and acetoclastic methanogenesis endmembers (Figure S3). We calculated ^D^*α*_p_ and ^D^*α*_s_ by the slopes, intercepts and *f*, following eqn. 7, and the carbon isotope fractionation factors from the isotope data of the ‘pure’ methylotrophic and acetoclastic methanogenesis.

The free parameters include ^13CD^*γ*_p_, ^13CD^*γ*_s_, ^DD^*γ*_p_, ^DD^*γ*_s_. The isotopic signatures of the ‘pure’ endmembers are used to get the best-fit values of these free parameters in the model. The values and 1-σ uncertainties of all input parameters are listed in Table 3.

To evaluate the effect of uncertainties on the modeled results, we conducted a Monte Carlo error propagation (MCEP) with the input values and uncertainties. Each parameter in the model is assumed to have a normal distribution with the measured or assigned mean value and 1-σ uncertainty. For each of the two methanogenic pathways, we ran the model using *δ*D_H2O_ as an independent variable, ranging from -150 to +9000 ‰ with a 50 ‰ increment. At each *δ*D_H2O_, we ran 1000 simulations and calculated the mean and standard deviations of the isotopic results (Figure 5).

## Acknowledgments

We thank Daniel Stolper for the contributions to the isotopic measurements of substrates. We thank Jonathan Gropp and Jiarui Liu for their in-depth discussions on the interpretation of the data. We also thank William Metcalf for offering the microbial strain used in this study.

## Funding

Simons Foundation Award 62388 (WDL)

NASA Exobiology 80NSSC21K0477 (EDY, WDL)

NASA Exobiology grant 80NSSC21K1240 (JFH)

Netherlands Organization for Scientific Research (N.W.O.)/Ministry of Education (OCW) grant SIAM 024002002 (SB, CUW, MJ)

ERC Synergy grant MARIX 854088 (MJ)

## Author contributions

Conceptualization: JLA, WDL.

Methodology: JL, AC, BCK, MAT, XF, GR, SB, EPM, AM, ALM, XF, JFH, WDL, RS.

Investigation: JL, JLA, AC, BCK, GR, MT, LG, SB, KM, YL, EPM, AM, ALM, XF, JFH, RS.

Visualization: JL, JLA.

Supervision: MF, JFH, CUW, MJ, EDY, WDL.

Funding: WDL, EDY, JFH, CUW, MJ.

Writing—original draft: JL, JLA.

Writing—review & editing: JL, JLA, BCK, GR, LG, SB, KM, YL, EPM, ALM, XF, JFH, AM, CUW, MJ, EDY, WDL.

## Data and materials availability

All data needed to evaluate the conclusions of this paper are available in the main text or the Supplementary Materials. The Python, MATLAB scripts and data frames used in the analysis are available on Figshare: https://doi.org/10.6084/m9.figshare.28011452.v1

## Supplementary Materials

**Figure S1:**
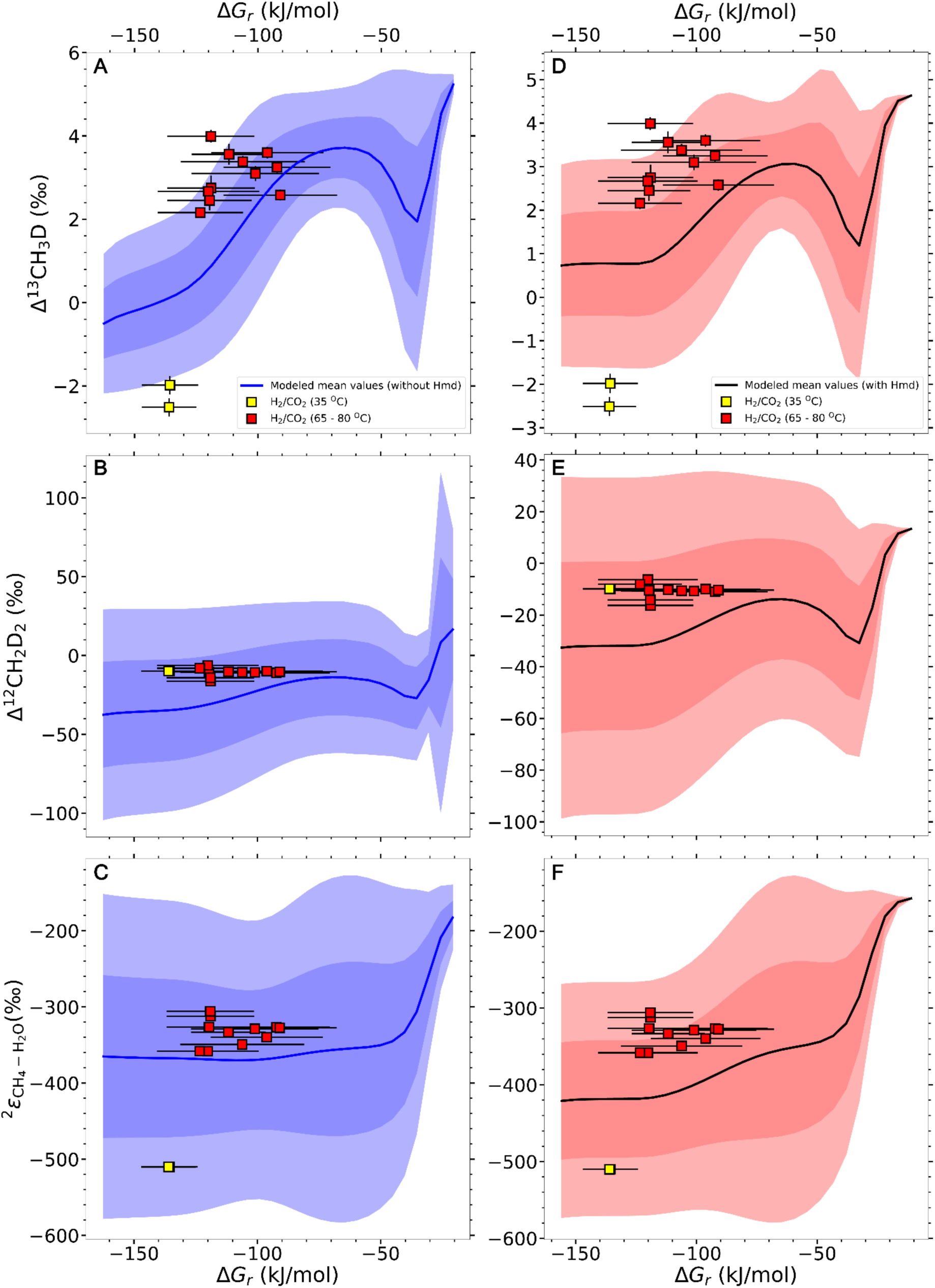
The variation of isotopic signatures with Gibbs free energy (ΔG_r_) of hydrogenotrophic methanogenesis. Panels A-C are the two clumped isotope signatures (Δ^13^CH_3_D, Δ^12^CH_2_D_2_) and hydrogen isotope fractionation between methane and water (*ε*_CH4-H2O_) using the model with H_2_-forming methylenetetrahydromethanopterin dehydrogenase (Hmd). Panels D-F are the modeled results without Hmd. The deep and shallow colored areas mark one and two standard deviations of the results, respectively. The hydrogenotrophic methanogenesis data in this study are shown in colored squares. The bars show the range of ΔG_r_ over the course of the experiments. The model is adopted from the previous study (*22*), and the uncertainties of the kinetic isotope effects (KIE) at each step are set as 1.5 times of the original uncertainties.

**Figure S2:**
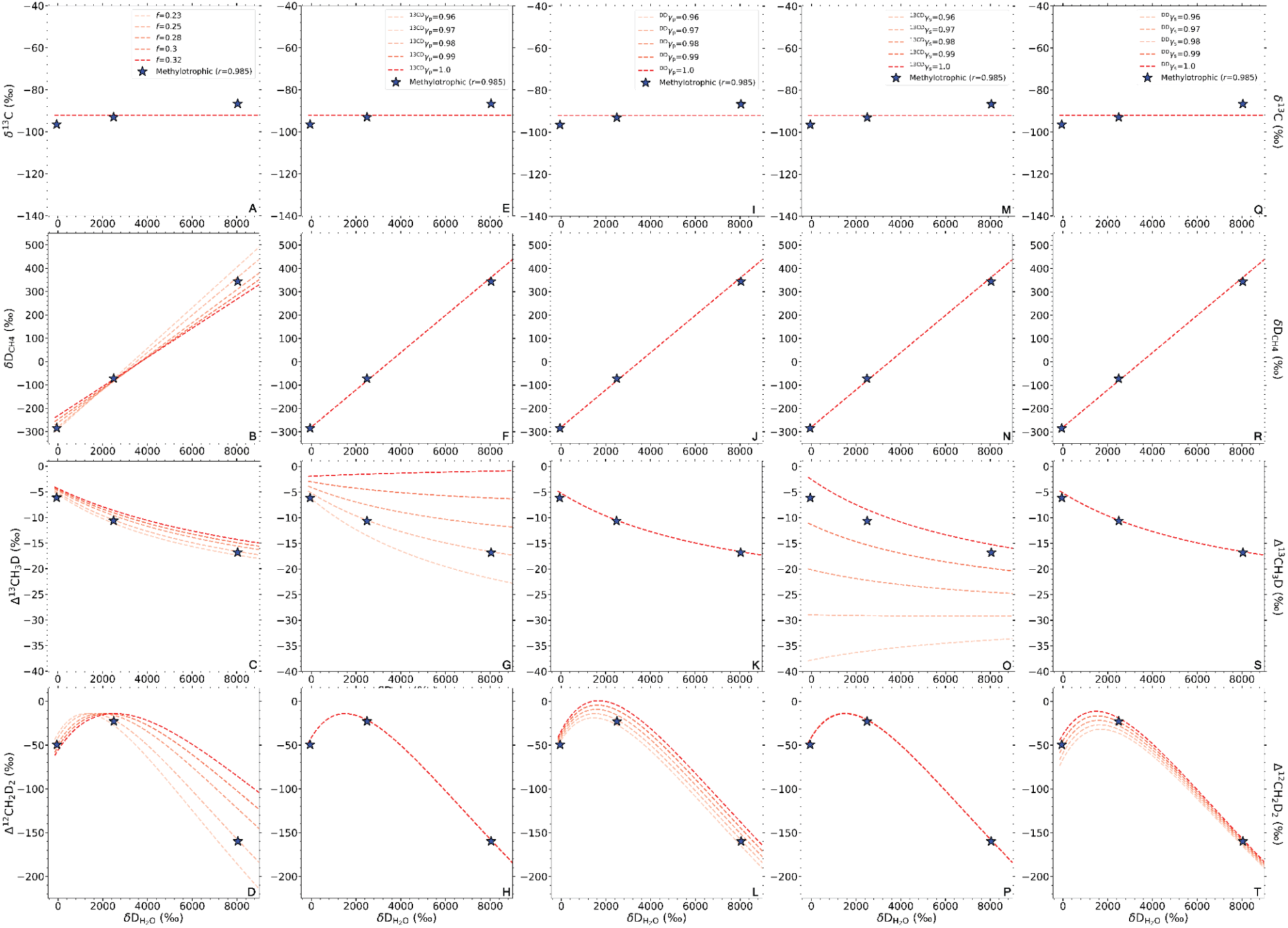
Sensitivity analysis of the five parameters (*f*, ^13CD^*γ*_p_, ^DD^*γ*_p_, ^13CD^*γ*_s_, ^DD^*γ*_s_) on the isotopic signatures in the combinatorial effect model. Panels A-D, E-H, I-L, M-P and Q-T show the effects of *f*, ^13CD^*γ*_p_, ^DD^*γ*_p_, ^13CD^*γ*_s_, ^DD^*γ*_s_ on the isotopic signatures (δ^13^C_CH4_, δD_CH4_, Δ^13^CH_3_D, Δ^12^CH_2_D_2_), respectively. Each parameter used in the sensitivity analysis is set at an initial value (see Table S4). At each test, one of the five parameters (*f*, ^13CD^*γ*_p_, ^DD^*γ*_p_, ^13CD^*γ*_s_, ^DD^*γ*_s_) changes within the values shown in the legends (also in Table S4). As a reference, the ‘pure’ methylotrophic methanogenesis endmembers are shown in stars.

**Figure S3:**
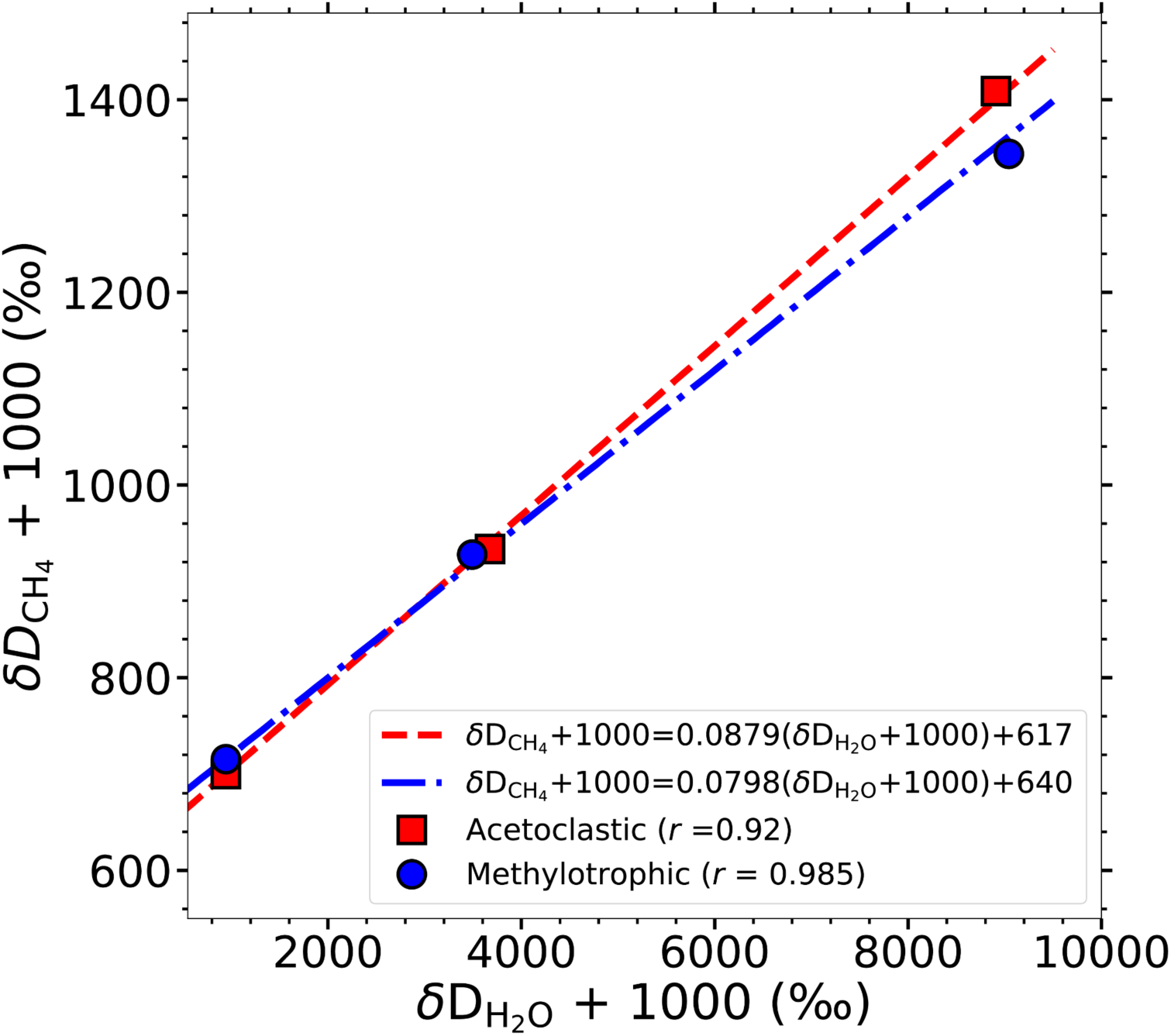
**Linear regression lines and equations between (δD_CH4_ + 1000) and (δD_H2O_ + 1000) for the ‘pure’ methylotrophic and acetoclastic endmembers.**

**Table S1:** Substrate isotope compositions.

| Substrate | $\delta^{13}\text{C}$ | 1-$\sigma^*$ | $\delta\text{D}^\dagger$ | 1-σ | $\Delta^{13}\text{CH}_3\text{D}$ | 1-σ | $\Delta^{12}\text{CH}_2\text{D}_2$ | 1-σ |
| --- | --- | --- | --- | --- | --- | --- | --- | --- |
| Radboud |  |  |  |  |  |  |  |  |
| Methanol | -37.57 | 1.23 | -65.03 | 0.32 | - | - | - | - |
| TMA | -41.67 | 0.34 | -109.73 | 0.16 | - | - | - | - |
| TMB | -37.59 | 1.32 | 1.97 | 0.19 | - | - | - | - |
| Dartmouth |  |  |  |  |  |  |  |  |
| CO<sub>2</sub> | -36.01 | 0.05 | - | - | - | - | - | - |
| Methanol | -41.34 | 0.02 | -54.11 | 0.25 | 0.842 | 0.352 | 6.26 | 3.78 |
| Acetate | -32.2 | 0.05 | -159.7 | 1.9 | - | - | - | - |

\* 1 standard deviations of the measurements
† The $\delta\text{D}$ values of methyl group/groups in the substrates.

**Table S2.** Net fractionation factors.

| Strain | Substrate | Temp | $^2\epsilon_{\text{CH}_4-\text{H}_2\text{O}}$ | 1-σ | $^{\text{D}}\alpha_{\text{CH}_4-\text{H}_2\text{O}}$ | 1-σ | $^{13}\alpha_{\text{CH}_4-\text{Carbon}}$ | 1-σ |
| --- | --- | --- | --- | --- | --- | --- | --- | --- |
| Rice/Radboud |  |  |  |  |  |  |  |  |
| *M. luminyensis* | CH<sub>3</sub>OH + H<sub>2</sub> | 37 °C | -232.78 | 0.05 | 0.77 | 0.00005 | 0.94 | 0.0012 |
| *M. luminyensis* | CH<sub>3</sub>OH + H<sub>2</sub> | 37 °C | -233.37 | 0.04 | 0.77 | 0.00004 | 0.94 | 0.0012 |
| *M. luminyensis* | TMA + H<sub>2</sub> | 37 °C | -325.04 | 0.04 | 0.67 | 0.00004 | 0.98 | 0.00034 |
| *M. luminyensis* | TMA + H<sub>2</sub> | 37 °C | -328.79 | 0.05 | 0.67 | 0.00005 | 0.97 | 0.00034 |
| *M. luminyensis* | TMA + H<sub>2</sub> | 37 °C | -328.06 | 0.04 | 0.67 | 0.00004 | 0.97 | 0.00034 |
| *M. shengliensis* | CH<sub>3</sub>OH | 65 °C | -237.2 | 0.05 | 0.76 | 0.00005 | 0.96 | 0.00122 |
| *M. shengliensis* | CH<sub>3</sub>OH | 65 °C | -231.72 | 0.05 | 0.77 | 0.00005 | 0.96 | 0.00123 |
| *M. shengliensis* | CH<sub>3</sub>OH | 65 °C | -230.79 | 0.05 | 0.77 | 0.00005 | 0.97 | 0.00124 |
| *M. shengliensis* | TMB | 65 °C | -244.01 | 0.05 | 0.76 | 0.00005 | 0.95 | 0.0013 |
| *M. mazei* | CH<sub>3</sub>OH | 39 °C | -249.35 | 0.05 | 0.75 | 0.00005 | 0.93 | 0.00118 |
| *M. mazei* | CH<sub>3</sub>OH | 39 °C | -249.22 | 0.05 | 0.75 | 0.00005 | 0.93 | 0.00118 |
| *M. mazei* | TMA | 39 °C | -346.65 | 0.04 | 0.65 | 0.00004 | 0.95 | 0.00033 |
| **Dartmouth, substrate** |  |  |  |  |  |  |  |  |
| *M. barkeri* | H<sub>2</sub>+CO<sub>2</sub> | 35 °C | -509.97 | 0.12 | 0.49 | 0.00012 | 0.96 | 0.00005 |
| *M. barkeri* | H<sub>2</sub>+CO<sub>2</sub> | 35 °C | -510.11 | 0.11 | 0.49 | 0.00011 | 0.96 | 0.00005 |
| *M. barkeri* | CH<sub>3</sub>OH | 35 °C | -254.17 | 0.2 | 0.75 | 0.0002 | 0.93 | 0.00002 |
| *M. barkeri* | CH<sub>3</sub>OH | 35 °C | -249.19 | 0.2 | 0.75 | 0.0002 | 0.94 | 0.00002 |
| *M. barkeri* | CH<sub>3</sub>OH | 35 °C | -237.47 | 0.2 | 0.76 | 0.0002 | 0.95 | 0.00002 |
| *M. barkeri* | Acetate | 35 °C | -278.03 | 0.31 | 0.72 | 0.00031 | 0.98 | 0.00005 |
| *M. barkeri* | Acetate | 35 °C | -278.98 | 0.31 | 0.72 | 0.00031 | 0.98 | 0.00006 |
| **UMass** |  |  |  |  |  |  |  |  |
| *M. jannaschii* | H<sub>2</sub> (full)+CO<sub>2</sub> | 80 °C | -312.21 | 0.14 | 0.69 | 0.00014 |  |  |
| *M. bathoardescens* | H<sub>2</sub> (full)+CO<sub>2</sub> | 80 °C | -326.42 | 0.12 | 0.67 | 0.00012 |  |  |
| *M. thermolithotrophicus* | H<sub>2</sub> (full)+CO<sub>2</sub> | 65 °C | -370.17 | 0.1 | 0.63 | 0.0001 |  |  |
| *M. thermolithotrophicus* | H<sub>2</sub> (full)+CO<sub>2</sub> | 65 °C | -358.14 | 0.09 | 0.64 | 0.00009 |  |  |
| *M. thermolithotrophicus* | H<sub>2</sub> (full)+CO<sub>2</sub> | 65 °C | -358.09 | 0.09 | 0.64 | 0.00009 |  |  |
| *M. jannaschii* | H<sub>2</sub> (full)+CO<sub>2</sub> | 80 °C | -305.84 | 0.14 | 0.69 | 0.00014 |  |  |
| *M. jannaschii* | H<sub>2</sub> (30 mL)+CO<sub>2</sub> | 80 °C | -333.32 | 0.14 | 0.67 | 0.00014 |  |  |
| *M. bathoardescens* | H<sub>2</sub> (30 mL)+CO<sub>2</sub> | 80 °C | -328.68 | 0.08 | 0.67 | 0.00008 |  |  |
| *M. thermolithotrophicus* | H<sub>2</sub> (30 mL)+CO<sub>2</sub> | 65 °C | -349.27 | 0.18 | 0.65 | 0.00018 |  |  |
| *M. jannaschii* | H<sub>2</sub> (10 mL)+CO<sub>2</sub> | 80 °C | -326.65 | 0.14 | 0.67 | 0.00014 |  |  |
| *M. thermolithotrophicus* | H<sub>2</sub> (10 mL)+CO<sub>2</sub> | 65 °C | -339.72 | 0.11 | 0.66 | 0.00011 |  |  |
| *M. bathoardescens* | H<sub>2</sub> (10 mL)+CO<sub>2</sub> | 80 °C | -327.5 | 0.11 | 0.67 | 0.00011 |  |  |

**Table S3.** The average isotope values of the ‘pure’ methylotrophic, acetoclastic, and methoxydotrophic endmembers. All data are shown in ‰.

| Substrates | $r$ | $\delta^{13}\text{C}$ | $\delta\text{D}$ | $\Delta^{13}\text{CH}_3\text{D}$ | $\Delta^{12}\text{CH}_2\text{D}_2$ |
| --- | --- | --- | --- | --- | --- |
| **Methanol** |  |  |  |  |  |
| -50 ‰ $\delta\text{D}_{\text{H}_2\text{O}}$ | 0.985 | -96.47 | -284.57 | -6.12 | -49.48 |
| +3000 ‰ $\delta\text{D}_{\text{H}_2\text{O}}$ | 0.985 | -93.01 | -72.19 | -10.62 | -22.77 |
| +8000 ‰ $\delta\text{D}_{\text{H}_2\text{O}}$ | 0.985 | -86.64 | 343.77 | -16.81 | -159.82 |
| **Acetate** |  |  |  |  |  |
| -50 ‰ $\delta\text{D}_{\text{H}_2\text{O}}$ | 0.92 | -46.06 | -299.99 | -2.67 | -49.65 |
| +3000 ‰ $\delta\text{D}_{\text{H}_2\text{O}}$ | 0.92 | -51.53 | -66.44 | -2.60 | -10.73 |
| +8000 ‰ $\delta\text{D}_{\text{H}_2\text{O}}$ | 0.92 | -52.17 | 408.97 | -3.13 | -198.75 |
| **TMB** |  |  |  |  |  |
| -50 ‰ $\delta\text{D}_{\text{H}_2\text{O}}$ | 0.66 | -102.92 | -237.42 | 2.30 | -52.50 |

**Table S4.** Values of the parameters used in the sensitivity test.

| Input parameters | Initial values | Tested values* |
| --- | --- | --- |
| **Fixed parameters** |  |  |
| Slope | 0.0798 | - |
| Intercept | 640.373 | - |
| $\delta D_{\text{methyl}}$ (‰) | -54.113 | - |
| $\delta^{13}\text{C}_{\text{methyl}}$ (‰) | -41.343 | - |
| $\Delta^{13}\text{CH}_2D_{\text{methyl}}$ (‰) | +0.842 | - |
| $\Delta^{12}\text{CD}_2\text{H}_{\text{methyl}}$ (‰) | +6.255 | - |
| $^{13}\alpha$ | 0.9471 | - |
| $r$ | 0.985 | - |
| $\delta D_{\text{H}_2\text{O}}$ (‰)<sup>†</sup> | [-150, 9000] | - |
| **Tested parameters** |  |  |
| $f$ | 0.25 | [0.23,0.25,0.28,0.3,0.32] |
| $^D\alpha_p^{\ddagger}$ | Slope/$f$ | - |
| $^D\alpha_s^{\ddagger}$ | Intercept/$((1-f)(\delta D_{\text{methyl}} + 1000))$ | - |
| $^{13}\text{CD}\gamma_p$ | 0.9700 | [0.96,0.97,0.98,0.99,1.00] |
| $^{\text{DD}}\gamma_p$ | 0.9700 | [0.96,0.97,0.98,0.99,1.00] |
| $^{13}\text{CD}\gamma_s$ | 0.9970 | [0.96,0.97,0.98,0.99,1.00] |
| $^{\text{DD}}\gamma_s$ | 0.9950 | [0.96,0.97,0.98,0.99,1.00] |

\* This shows the list of values used in the sensitivity test. During the test of each parameter, the values are chosen from the list, while the rest are set at the initial values. <sup>†</sup> Similar to the model, the $\delta D_{\text{H}_2\text{O}}$ in each test ranges from -150 to 9000 ‰, with a 50 ‰ increment. <sup>‡</sup> $^D\alpha_p$ and $^D\alpha_s$ vary with $f$ in the test.

| Reagent | Amount per L distilled H<sub>2</sub>O |
| --- | --- |
| K<sub>2</sub>HPO<sub>4</sub> | 0.348 g |
| KH<sub>2</sub>PO<sub>4</sub> | 0.227 g |
| NH<sub>4</sub>Cl | 0.5 g |
| MgSO<sub>4</sub> x 7H<sub>2</sub>O | 0.5 g |
| CaCl<sub>2</sub> x 2H<sub>2</sub>O | 0.25 g |
| NaCl | 2.25 g |
| NaHCO<sub>3</sub> | 0.85 g |
| Vitamin solution 1000x | 1 mL |
| SL – 6 (altered for trace element content)* | 1 mL |
| Yeast extract | 2 g |
| Casiton | 2 g |
| NaCH<sub>3</sub>COO | 1 g |
| Resazurin (1mg/mL) | 1 mL |

\*1000 × trace element stock solution: 1.35 g/L FeCl<sub>2</sub> × 4 H<sub>2</sub>O, 0.1 g/L MnCl<sub>2</sub> × 4 H<sub>2</sub>O, 0.024 g/L CoCl<sub>2</sub> × 6 H<sub>2</sub>O, 0.1 g/L CaCl<sub>2</sub> × 2 H<sub>2</sub>O, 0.1 g/L ZnCl<sub>2</sub>, 0.025 g/L CuCl<sub>2</sub> × 2 H<sub>2</sub>O, 0.01 g/L H<sub>3</sub>BO<sub>3</sub>, 0.024 g/L Na<sub>2</sub>MoO<sub>4</sub> × 2 H<sub>2</sub>O, 0.22 g/L NiCl<sub>2</sub> × 6 H<sub>2</sub>O, 0.017 g/L Na<sub>2</sub>SeO<sub>3</sub>, 0.004 g/L Na<sub>2</sub>WO<sub>4</sub> × 2 H<sub>2</sub>O, 12.8 g/L nitrilotriacetic acid.

| Reagent | Amount per L distilled H<sub>2</sub>O |
| --- | --- |
| KCl | 0.34 g |
| NH<sub>4</sub>Cl | 0.25 g |
| K<sub>2</sub>HPO<sub>4</sub> | 0.20 g |
| NaCl | 24 g |
| MgCl<sub>2</sub> x 6H<sub>2</sub>O | 10.2 g |
| Yeast extract | 2 g |
| Resazurin | 1 mL of 0.1% solution |
| NaHCO<sub>3</sub> | 2.5 g |
| Vitamin solution 1000x | 1 mL |
| SL – 6 (altered for trace element content)* | 1 mL |

\*1000 × trace element stock solution: 1.35 g/L FeCl<sub>2</sub> × 4 H<sub>2</sub>O, 0.1 g/L MnCl<sub>2</sub> × 4 H<sub>2</sub>O, 0.024 g/L CoCl<sub>2</sub> × 6 H<sub>2</sub>O, 0.1 g/L CaCl<sub>2</sub> × 2 H<sub>2</sub>O, 0.1 g/L ZnCl<sub>2</sub>, 0.025 g/L CuCl<sub>2</sub> × 2 H<sub>2</sub>O, 0.01 g/L H<sub>3</sub>BO<sub>3</sub>, 0.024 g/L Na<sub>2</sub>MoO<sub>4</sub> × 2 H<sub>2</sub>O, 0.22 g/L NiCl<sub>2</sub> × 6 H<sub>2</sub>O, 0.017 g/L Na<sub>2</sub>SeO<sub>3</sub>, 0.004 g/L Na<sub>2</sub>WO<sub>4</sub> × 2 H<sub>2</sub>O, 12.8 g/L nitrilotriacetic acid.

| Component | Amount | MW | final concentration |
| --- | --- | --- | --- |
| NaCl | 23.4 g | 58.45 | 400 mM |
| NaHCO<sub>3</sub> | 3.8 g | 84.01 | 45 mM |
| KCl | 1.0 g | 74.56 | 13 mM |
| 0.1% resazurin | 1.0 mL | 251.2 | 4 μM |
| Trace elements | 10.0 mL | - | see formulation |
| Vitamin solution | 10.0 mL | - | see formulation |

| Component | Amount | MW | final concentration |
| --- | --- | --- | --- |
| MgCl<sub>2</sub>.6H<sub>2</sub>O | 11.0 g | 203.31 | 54 mM |
| CaCl<sub>2</sub>.2H<sub>2</sub>O | 0.3 g | 147.02 | 2 mM |

| Component | Amount | MW | final concentration |
| --- | --- | --- | --- |
| 1.0 M KH<sub>2</sub>PO<sub>4</sub> | 5.0 mL | - | 5 mM |
| NH<sub>4</sub>Cl | 1.0 g | 53.49 | 19 mM |
| Cysteine.HCl | 0.5 g | 175.6 | 2.8 mM |
| 0.2 M Na<sub>2</sub>S.9H<sub>2</sub>O | 2.0 mL | ----- | 0.4 mM |
| Anaerobic, pH=6.8 |  |  |  |

Optional Components (all from sterile anaerobic stock solutions)
| Component | Amount | MW | final concentration |
| --- | --- | --- | --- |
| 100 % Methanol | 5.0 mL/L | - | 125 mM |
| 5 M TMA-HCl | 10.0 mL/L | - | 50 mM |
| 20 % yeast extract | 10.0 mL/L | - | 0.2 % |
| 4.0 M Na acetate | 10.0 mL/L | - | 40 mM |
| 10 % So Slurry | 0.2 mL/20 mL | - | 0.1 % |
| 40 % glucose | 0.8 mL/20 mL | - | 1.0 % |

Modified Trace Elements Solution
| Compound | Amt./Liter | MW | Conc. | Media Conc. |
| --- | --- | --- | --- | --- |
| Nitrilotriacetic acid<br>(Trisodium salt) | 1.5 g | 257.1 | 5.8 mM | 58 µM |
| Fe(NH<sub>4</sub>)<sub>2</sub>(SO<sub>4</sub>)<sub>2</sub> | 0.8 g | 392.1 | 2 mM | 20 µM |
| Na<sub>2</sub>SeO<sub>3</sub> | 0.2 g | 172.9 | 1.1 mM | 11 µM |
| CoCl<sub>2</sub>.6H<sub>2</sub>O | 0.1 g | 237.9 | 0.4 mM | 4 µM |
| MnSO<sub>4</sub>.H<sub>2</sub>O | 0.1 g | 169 | 0.6 mM | 6 µM |
| Na<sub>2</sub>MoO<sub>4</sub>.2H<sub>2</sub>O | 0.1 g | 241.9 | 0.4 mM | 4 µM |
| Na<sub>2</sub>WO<sub>4</sub>.2H<sub>2</sub>O | 0.1 g | 329.9 | 0.3 mM | 3 µM |
| ZnSO<sub>4</sub>.7H<sub>2</sub>O | 0.1 g | 287.5 | 0.3 mM | 3 µM |
| NiCl<sub>2</sub>.6H<sub>2</sub>O | 0.1 g | 237.7 | 0.4 mM | 4 µM |
| H<sub>3</sub>BO<sub>3</sub> | 0.01 g | 61.83 | 0.16 mM | 1.6 μM |
| CuSO<sub>4</sub>·5H<sub>2</sub>O | 0.01 g | 249.7 | 40 μM | 0.4 μM |

Vitamin Solution
| Compound | Amt./Liter | MW | Conc. | Media Conc. |
| --- | --- | --- | --- | --- |
| *p*-Aminobenzoic acid | 10 mg | 137.1 | 73 μM | 729 nM |
| Nicotinic acid | 10 mg | 123.1 | 81 μM | 812 nM |
| Ca pantothenate | 10 mg | 238.3 | 42 μM | 419 nM |
| Pyridoxine HCl | 10 mg | 205.6 | 49 μM | 486 nM |
| Ribflavin | 10 mg | 376.4 | 27 μM | 266 nM |
| Thiamine HCl | 10 mg | 337.3 | 30 μM | 296 nM |
| Biotin | 5 mg | 244.3 | 20 μM | 204 nM |
| Folic acid | 5 mg | 441.4 | 11 μM | 113 nM |
| α-Lipoic acid | 5 mg | 206.3 | 24 μM | 242 nM |
| Vitamin B12 | 5 mg | 1355.4 | 3.7 μM | 37 nM |

Sterilize by filtration using a 0.22 micron filter
Store in the dark at 4 °C.

### UMass

Each 60 mL serum bottle contains 25 mL of DSM 282 growth medium which contains (per liter): 780 mL of distilled deionized H_2_O, 200 mL 5× salts (containing 150 g of NaCl, 20.5 g of MgCl_2_•6H_2_O, 17 g of MgSO_4_•7H_2_O, 1.65 g of KCl, 1.25 g of NH_4_Cl, 0.7 g of CaCl_2_•2H_2_O, 0.7 g of K_2_HPO_4_), 10 mL of DSM 141 trace minerals solution, 10 mL of DSM 141 vitamin solution, 1 g of NaHCO_3_, 1 g of Na_2_s_2_O_3_, 0.10 ml of 0.1% (Na_2_WO_4_•2H_2_O, 0.1% Na_2_SeO_3_) and 0.05 mL of 0.5% (w/v) resazurin solution. The medium was pH balanced to pH 6.00 ± 0.05 using 1 M HCl. Before inoculation, each bottle was reduced with 0.25 ml cysteine-HCl and 0.25 ml Na_2_S•9H_2_O.

### Estimation of ΔG_r_ in hydrogenotrophic methanogenesis

The net Gibbs free energy yields (ΔG_r_) and the corresponding isotopic signatures in hydrogenotrophic methanogenesis experiments are estimated following the model described in the previous study (*22*). The net Gibbs free energy is calculated using the standard state Gibbs free energy (ΔG_0_), as well as dissolved concentrations of H_2_, CO_2_ and CH_4_:

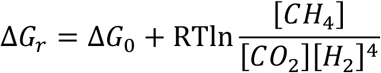

Where the bracketed values are dissolved gas concentrations in mol/L. The standard Gibbs free energy at the given temperature is calculated from the Van’t Hoff equation, following the previous literature (*80*). The dissolved H_2_, CO_2_ and CH_4_ concentrations in the ΔG_r_ calculations are obtained from Henry’s law, using the headspace gas concentrations. The Henry’s law constants for H_2_, CO_2_ and CH_4_ are 7.5, 1.5 and 3.3 µmol/(m^3^*Pa), respectively (*81*). The initial H_2_, CO_2_ partial pressures are calculated from the initial growth conditions described in the Materials and Methods Section, while the initial CH_4_ is set at a very low level (1/1000 of the initial CO_2_ partial pressures). We estimated the partial pressures of the gasses at the end of the experiments by calculating the H_2_ and CO_2_ consumptions from CH_4_ production, following the stoichiometry of the methanogenic reaction:

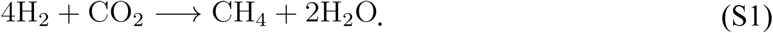

We ran the model developed by Gropp et al. (*22*) with *δ*^13^C and *δ*D values of the substrates used in this study (-36.01 and -50.00 ‰, respectively). We also assigned larger uncertainties (1.5 times the original value) for the kinetic isotope effects of each reaction step in the model, since large uncertainties remain in those values (*22*). The modeled results, both with Hmd and without Hmd activities, are shown in Figure S1.

### Calculation of methanol consumption by stoichiometry

In methylotrophic methanogenesis, methanol is disproportionated into CH_4_ and CO_2_:

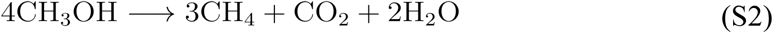

Therefore, we can calculate the consumption of methanol from the production of methane (Table 1) by stoichiometry. The consumption of methanol during the cultivation at Dartmouth College ranges from 181 to 473 µmol, corresponding to 14 to 38 % of the initial methanol (1250 µmol of initial methanol).

### Calculation of mixing

The mixing between the hydrogenotrophic and acetoclastic/methylotrophic endmembers are calculated using the ratios of the isotopes or isotopologues. For bulk isotopes (*δ*^13^C and *δ*D), the *δ* values are first converted to fractional abundance (*F*):

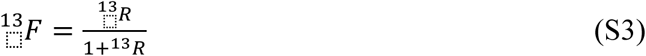

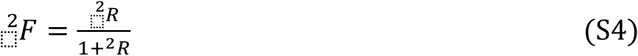

Where *R* is the ratio of heavy to light isotopes (D/H or ^13^C/^12^C) derived from the *δ* values. For the clumped isotopologues (Δ^13^CH_3_D and Δ^12^CH_2_D_2_), the absolute abundances of isotopologues that are measured on the instrument are used, which are calculated from the absolute abundances of isotopologues in the reference gas on Panorama (*67*). The mixing effect is then calculated as:

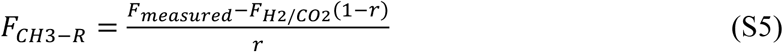

Or

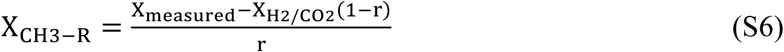

Where “CH3-R”, “measured”, and “H2/CO2” in the subscripts denote the ‘pure’ methylotrophic or acetoclastic methanogenesis, measured samples, and hydrogenotrophic methanogenesis by *M. barkeri*, respectively. *F* and *X* are the fractional or absolute abundances of heavy isotopes or clumped isotopologues. *r* is the portion of methane derived from the methylotrophic or acetoclastic methanogenesis in the mixture. The calculated fractional abundances or absolute abundances are then converted to *δ*^13^C, *δ*D, Δ^13^CH_3_D and Δ^12^CH_2_D_2_ using eqn. 1 to 4 in the main text. Specifically, the stochastic distributions in eqn. 3 and 4 are calculated from the random distribution of isotopes (*13*). The derived ‘pure’ endmembers are listed in Table S3.

